# Cat_Wiz: A stereochemistry-guided toolkit for locating, diagnosing and annotating Mg^2+^ ions in RNA structures

**DOI:** 10.1101/2025.10.04.677962

**Authors:** Nawavi Naleem, Anja Henning-Knechtel, Serdal Kirmizialtin, Pascal Auffinger

## Abstract

Misassigned Mg^2+^ ions are pervasive in RNA structural databases, obscuring mechanistic interpretation, undermining comparative analyses and compromising machine-learning training sets. Here, we present *Cat_Wiz*, a *Coot*-integrated, stereochemistry guided toolkit that facilitates the localization, diagnosis, correction and annotation of Mg^2+^ binding sites. *Cat_Wiz* comprises three modules: *MG_diagnosis which* validates and regularizes existing assignments; *MG_detect which* identifies unmodelled ion binding sites; and *MG_clamp which* classifies recurrent Mg^2+^ clamp motifs. *Cat_Wiz* also includes a complete binding site annotation system. The stereochemical principles implemented in *Cat_Wiz* were derived from an earlier analysis of the 1.55 Å resolution *Escherichia coli* ribosome and from surveys of the *Cambridge Structural Database.* These principles provide a robust experimental foundation for characterizing Mg^2+^ binding sites. Applications to ribosomes, hammerhead ribozymes, group I introns, and quaternary RNA assemblies demonstrate that *Cat_Wiz* rapidly locates overlooked ions, corrects misassignments, and improves stereochemical fidelity in hours rather than days. Beyond refinement, *Cat_Wiz* generates curated data that can seed diverse machine-learning and AI models. This transparent, cost-effective framework establishes reproducible standards for RNA-ion assignments and will drive progress in the design of RNA 3D architectures through the identification of unique Mg^2+^-dependent backbone folds. *Cat_Wiz* is also applicable to Mg^2+^ binding sites in proteins and all related biomolecular systems since it is based on universal stereochemical principles.

## INTRODUCTION

Hexahydrated and partially dehydrated magnesium (Mg^2+^) ions are central to RNA structure, folding, and catalysis (1–7). Accurate placement and subsequent validation of Mg^2+^ ions in experimental models are therefore critical for mechanistic interpretation, comparative mining and other computational use. While several studies have investigated Mg^2+^ coordination to RNA (5,8–14), misassignments remain common in deposited models where ions are often placed in environments incompatible with their characteristic octahedral geometry, thereby compromising database surveys and associated inferences (15–18).

In 2022, a cryo-EM structure of a 70S *Escherichia coli* ribosome was solved at 1.55 Å (PDBid: **8b0x**). This high-resolution structure provided an excellent opportunity to codify Mg^2+^ binding principles that prompted the creation of a robust experimentally based coordination pattern index (14,19). Consequently, the **8b0x** model was revised to correct Mg^2+^ assignments and missing water molecules mandatory to complete octahedral coordination shells. This process significantly increased the number of authenticated Mg^2+^ binding sites and revealed recurrent assignment pathologies.

Manual curation at such scales is particularly time-consuming, especially for high-resolution structures like ribosomes that can comprise over 20,000 solvent particles (14). Automated methods ranging from deep-learning classifiers to segmentation-guided solvent placement and molecular dynamics (MD) simulations remain vulnerable to local map heterogeneity and violation of well-established stereochemical boundaries (14,20–22). These drawbacks motivated the development of a transparent, user-friendly and stereochemistry-guided approach for routine Mg^2+^ assignment and validation of X-ray and cryo-EM models. The data gathered on Mg^2+^ binding motifs can further support the development of accurate RNA three-dimensional structure prediction tools (23–27).

Herein, we present *Cat_Wiz* (*“Cat”* stands for cation and *“Wiz”* for wizard), an open suite of Python scripts developed for the *Coot* molecular modeling package (28). *Cat_Wiz* comprises three modules (**Figure 1**): *MG_diagnosis* identifies assignment issues and provides an annotated summary of stereochemically validated Mg^2+^ binding sites while providing tools to complete and regularize Mg^2+^ coordination shells; *MG_detect* scans masked maps around coordinating O/N atoms in RNA (*MG_detect_RNA*) and proteins (*MG_detect_Prot*) to locate unmodelled Mg^2+^ ions; finally, *MG_clamp* identifies, classifies, and annotates Mg^2+^ clamps (see **Figure 2**), a recurrent motif in RNA systems (3,5,8,13,14,29–31).

**Figure 1.**
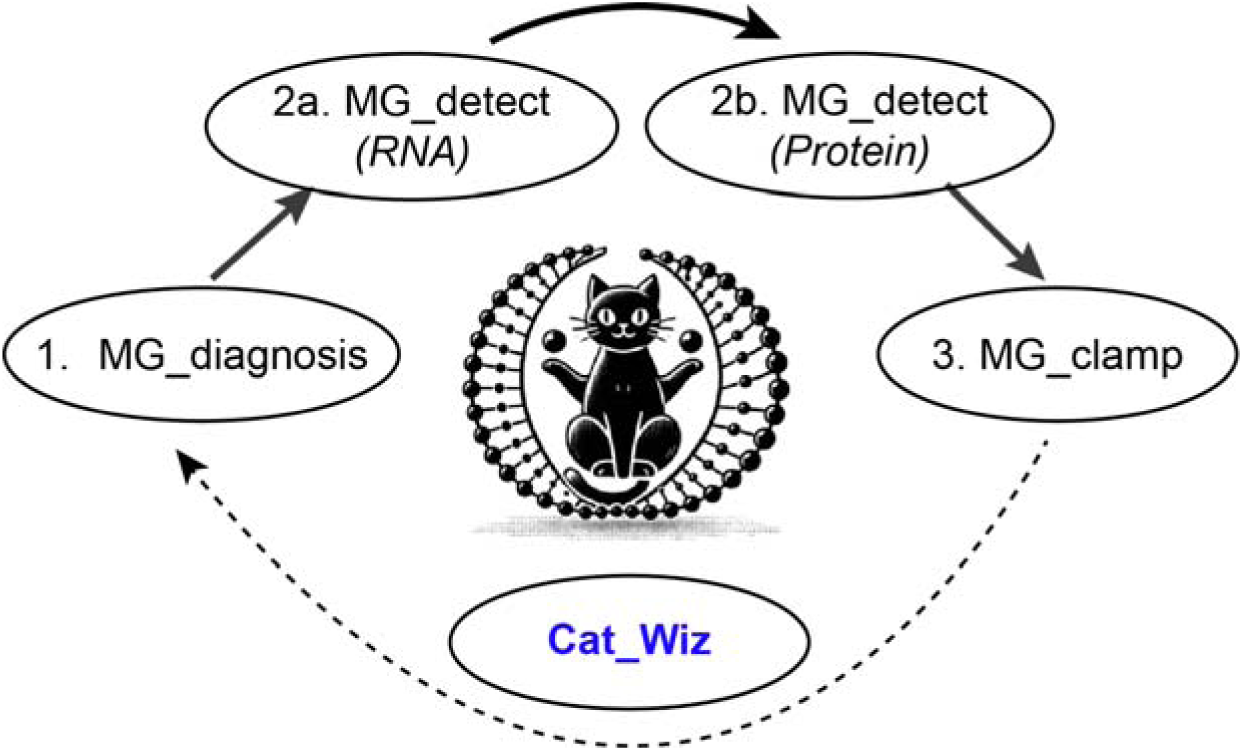
*Cat_Wiz* workflow. The diagram illustrates the recommended sequence for using the four independent Mg^2+^ related *Cat_Wiz* scripts. It is advised to re-run the *MG_diagnosis* script whenever structural changes were made to ensure the accuracy of the final model.

**Figure 2.**
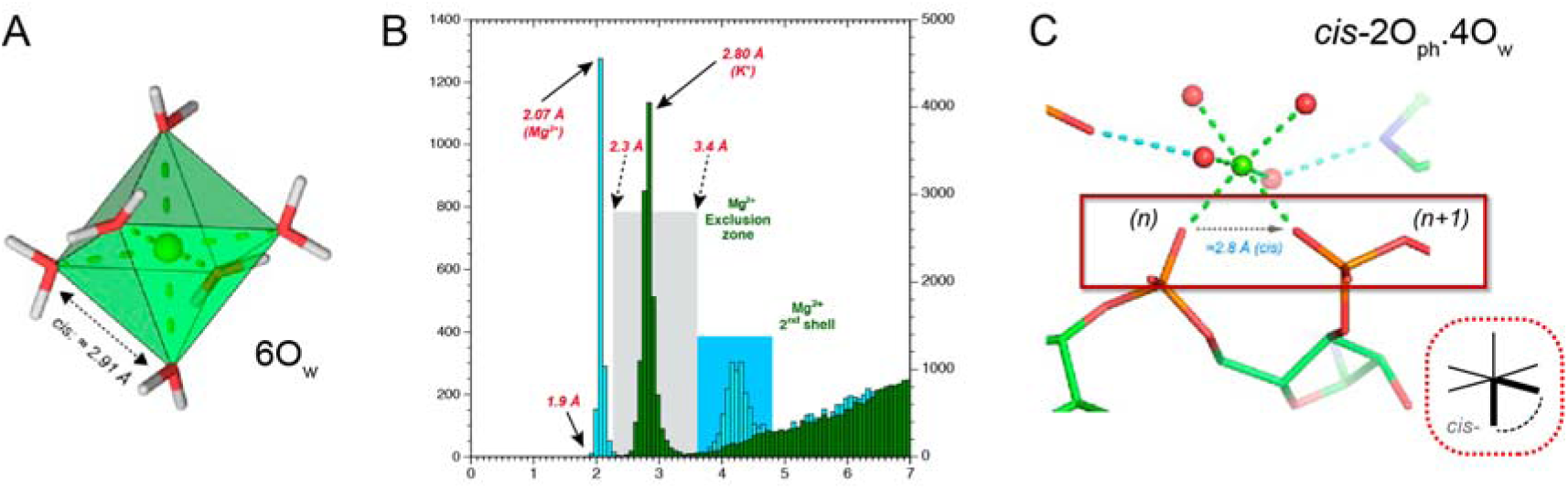
Stereochemical features of Mg^2+^ ions and Mg^2+^ clamps. (**A**) A hexahydrated Mg^2+^ ion (6O_w_) from a high-resolution *Cambridge Structural Database* (CSD) entry, showing characteristic *d*(O_w_…O_w_) distances around 2.9 Å (14). (**B**) Histograms of Mg^2+^ (cyan) and K^+^ (green) to oxygen coordination distance from the CSD. The distinct ∼0.7 Å distance between the two peaks is a reliable criterion for distinguishing these ions. The grey rectangle marks the Mg^2+^ exclusion zone, while the blue rectangle identifies its second coordination shell. (**C**) The geometry of a *cis-*2O_ph_.4O_w_ magnesium clamp involving two consecutive phosphate groups.

We demonstrate *Cat_Wiz* on four representative cases spanning different experimental modalities and resolutions: *(i)* the 1.55 Å resolution cryo-EM **8b0x** ribosome model (13), *(ii)* historically debated Mg^2+^ binding in hammerhead ribozymes (15,32), *(iii)* X-ray and near-atomic cryo-EM *Tetrahymena* group I intron structures, the latter having been used as a target in the 2024 CASP competition (26,33) and *(iv)* cryo-EM quaternary RNA assemblies with resolutions above 2.6 Å (34). Across all these cases, *Cat_Wiz* successfully rectified incorrect assignments, recovered missing hydrated ions and differentiated Mg^2+^ from K^+^ by enforcing representative coordination distances, thereby improving the fidelity of deposited structures. Additionally, *Cat_Wiz* can assist in the refinement and interpretation of nucleic acid and protein systems.

The stereochemical Mg^2+^ priors we defined in **Figure 2** were derived from small-molecule statistics through the *Cambridge Structural Database* (CSD) and described elsewhere (11,12,14,35). Several community resources already assist with metal-site identification or diagnosis. Among them, *CheckMyMetal* (CMM) performs geometry- and valence-based validation of metal-binding sites while providing an interactive platform applicable to X-ray and cryo-EM models (36–44). In this context, CMM and *Cat_Wiz* are complementary tools.

*Cat_Wiz* also complements popular refinement programs such as REFMAC5 and Phenix (45,46) by providing stereochemically-aware Mg^2+^ binding site evaluation. The suite offers a set of restraints to stabilize Mg^2+^ octahedra in lower-resolution regions. By embedding prior knowledge rather than learning from noisy structures, *Cat_Wiz* provides a fast, transparent refinement solution for solvent particles and Mg^2+^ ions that can enhance machine learning (ML) and artificial intelligence (AI) workflows. In the following sections, we *(i)* outline the current *Cat_Wiz* stereochemical nomenclature in detail, *(ii)* describe the different *Cat_Wiz* modules, *(iii)* apply the toolkit to the case studies introduced above and *(iv)* address *Cat_Wiz* limitations and future extensions.

## METHODS

### *Cat_Wiz* installation in *Coot*

*Cat_Wiz* is a suite of independent Python scripts (**Figure 1**) developed for the molecular modeling program *Coot* (28). To use the scripts, a coordinate file (PDB or mmcif) and a corresponding map file derived from X-ray or cryo-EM experiments must first be loaded into Coot. The user can then navigate to the *“Calculate”* menu, select *“Run scripts”,* and *choose* the desired script from the local directory. Upon first use, particularly with large structures such as ribosomes, *Cat_Wiz* requires a few minutes to generate a map smoothed by a factor of 1.25 (default value). During this process, the *Coot* console displays *“waiting”* messages. For X-ray structures, structure factor files (.sf) provided by the PDB must be converted into an *.mtz* file using the SF-tool web server (https://sf-tool.wwpdb.org) (47). Default cutoff parameters were empirically derived and can be adjusted in the setup header of the scripts that were tested with *Coot* version 0.9.8.98. A version adapted for *Coot* 1.xxx is currently under development. For additional guidance, users are referred to the updated help pages provided with each module. It is advised to exercise caution when working with *“over”-*sharpened maps that might alter results of the *MG_detect* module and occasionally lead to zero or negative *B-factors* at the refinement stage in Phenix. Next, we first describe additions to a previously reported Mg^2+^ binding site nomenclature (13) before detailing the specific features of each module.

### Mg^2+^ binding site nomenclature description and additions

*Cat_Wiz* is based on a Mg^2+^ binding site nomenclature derived from the MgRNA study (10–12,14) and summarized in (**Figure 3**). This nomenclature reflects the strict octahedral coordination geometry of Mg^2+^ characterized by *d*(Mg^2+^…O/N) coordination distances of 2.07/2.19 Å, with the slightly longer distance for corresponding nitrogen coordination. The system uses *cis-/trans-/fac-/mer-/CIS-/TRANS*-prefixes together with the following atom type identifiers: *“O_ph_”* for OP1/OP2 phosphate anionic oxygens; *“O_r_”* for O2’/O4’ ribose oxygens and O3’/O5’ phosphate oxygens; *“O_b_”* for O2/O4/O6 nucleobase oxygens; and *“N_b_”* for non-protonated N1/N3/N7 nucleobase nitrogens.

- *General nomenclature:* As an example, 2O_ph_.4O_w_ denotes a hexacoordinated ion bound to two O_ph_ atoms and four water molecules (**Figure 2C**). The prefixes *cis-* and *trans-* describe the relative orientation of the two O_ph_ atoms. When three non-water atoms adopt a *cis-* arrangement, the corresponding isoforms is noted *fac-*3O_ph_.3O_w_ (*facial*). When one O_ph_ pair is in *trans-* (10,14) configuration, the site is named *mer-*3O_ph_.3O_w_ (*meridional)*. When four non-water atoms coordinate the ion, as in 4O_ph_.2O_w_, the orientation of the two water molecules defines the *cis-/trans-* terminology. To distinguish these binding types from the more common *cis-/trans-*2O_ph_.4O_w_ types, the capital prefixes *CIS-/TRANS-* are used. This nomenclature can be extended to any combination of O_ph_/O_b_/O_r_/N_b_ atoms. For proteins, additional atom types are defined, i.e., O_bb_ for amino acid backbone oxygens (replacing the previously used O_back_ denomination (10,14)), O_coo_ for carboxyl oxygens (Asp/Glu), O_cno_ for carbonyl oxygens (Asn/Gln), O_coh_ for hydroxyl oxygens (Thr/Ser/Tyr), and N_his_ for non-protonated histidine nitrogens (ND1/NE2).
- *Water mediated Mg^2+^ clamps:* In rare instances, interactions between O/N atoms separated by < 3.4 Å are stabilized by both, a direct Mg^2+^ contact and a water-mediated interaction (**Figure 3**). Hence, the corresponding O_ph_.5O_w_ or O_b_.5O_w_ magnesium binding type become *“water-mediated”* clamps. We previously hypothesized that some of these patterns may represent precursors to more standard clamp geometries (14). Due to structural compaction, these water-mediated contacts also appear in more complex motifs but are not specifically annotated by the *Cat_Wiz* modules for the sake of nomenclature simplification.
- *Ribose O3’/O5’ atoms and Mg^2+^ coordination:* Recent investigations on group I and II introns have described catalytic sites with suspected Mg^2+^…O3’ contacts (48,49). This observation is puzzling as direct coordination of divalent ions to O3’ atoms have not been observed in the ensemble of structures we analyzed. Therefore, such contacts are not included in **Figure 3**. The only exception identified to date involves terminal nucleotides where the O3’-H group can coordinate to a Mg^2+^ ion, either alone or in combination with the O2’-H of the same ribose (50).
- *Monohydrated Mg^2+^ ions:* Binding patterns involving five non-water molecules are uncommon (**Figure 3**). One such site was modelled in the X-ray structure of *Oceanobacillus iheyensis* group II intron (PDBid: **5j01**) at 3.39 Å resolution (50). In this case, the ion coordinates to the 2’-OH and 3’-OH hydroxyl groups of a 3’-terminal uridine near the catalytic site. The organization of this motif may follow distinct binding rules related to two terminal-residue hydroxyl groups and may contribute to stabilization of leaving groups during catalysis.
- *Non-hydrated Mg^2+^ ions:* Coordination of six non-water atoms has not yet been observed in nucleic acid structures, likely due to steric constraints imposed by phosphate and nucleobase groups (**Figure 3**). This coordination mode has, however, been observed in a protein structure in which the six oxygen atoms belong to protein carbonyl and carboxylate groups as well as to three phosphate groups from a 2’-deoxyguanosine-5’-triphosphate (DGT) ligand (PDBid: **8ef9**) (51). Both monohydrated and fully dehydrated patterns are shown in grey in **Figure 3** as they have not been observed in representative nucleic acid systems to date.
- Mg^2+^ *coordination to metabolites, drugs and buffer molecules:* Examples include coordination to the tetracycline antibiotic in a ribosome structure (TAC; PDBid: **8cgj**) (52), to ZMP in a riboswitch (PDBid: **4xwf**) (53), or more commonly to di- and triphosphate metabolites (5). Mg^2+^ can also bind to oxygen atoms from buffer components like (4S)-2-methyl-2,4-pentanediol (MPD; PDBid: **7mky**) (54) or acetates (ACT/ACY; PDBid: **1yvp**) (55). Occasionally, non-oxygen atoms such as fluoride, form polyatomic clusters with Mg^2+^, as observed in fluoride riboswitches (PDBid: **4enc**) (56). Direct coordination of Mg^2+^ to a cacodylate anion (CAC: AsO_2_(CH_3_)_2_^−^; PDBid: **4xvf**) has also been reported.
- *Ligand nomenclature:* Currently, the *Cat_Wiz* scripts account for most modified nucleotides but do not consider Mg^2+^ binding to ligands. For describing such sites, we suggest using the O_L_ and N_L_ atom-type identifiers. For example, the tetracycline-bound ion in PDB entry **8cgj** would be described as a *TRANS-*2O_b_.2O_L_.2O_w_ or, more specifically, *TRANS-*2O_b_.2O_TAC_.2O_w_ when using the PDB ligand code for tetracycline (TAC). This example illustrates the versatility and extensibility of the Mg^2+^ annotation code.

**Figure 3.**
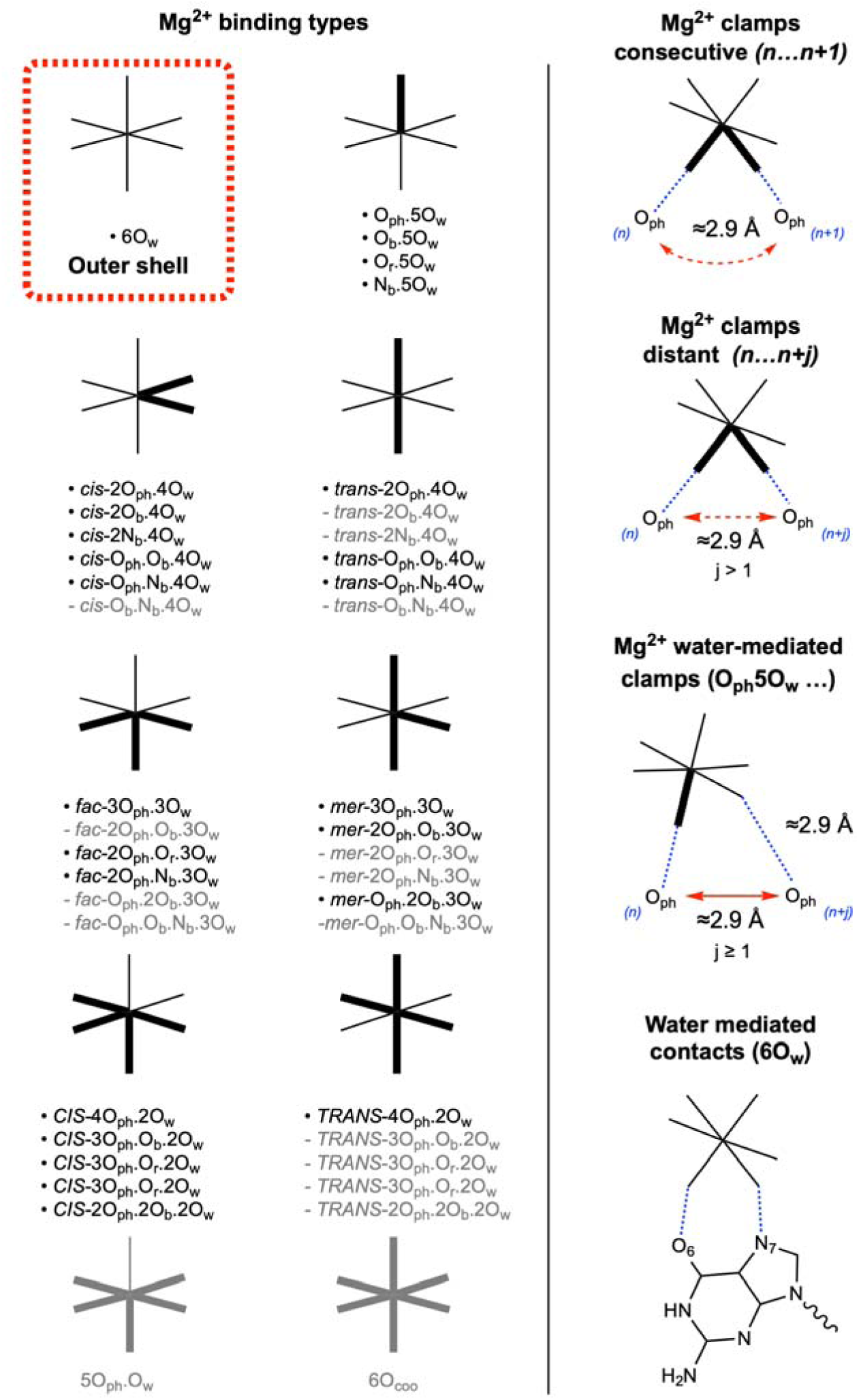
Updated view of Mg^2+^ binding motifs. (Left) This panel presents all inner- and outer-shell Mg^2+^ binding motifs identified in **9q87** and is an update of the list derived from **8b0x** (14). For clarity, motifs involving ribosomal proteins have been excluded. Greyed out binding types are until now associated with weak, insufficient, or no structural evidence. Binding to O3’ occurs only in exceptional configurations involving terminal residues and is not mentioned in the figure. Binding to O4’/O5’ has not been observed. (Right) The top two figures illustrate a classic Mg^2+^ clamp, in which the ion coordinates in *cis-* to two phosphate atoms, either through consecutive (j=1) or distant nucleotides (j>1). The third figure depicts a water-mediated O_ph_.5O_w_ motif. The bottom figure shows an outer-shell 6O_w_ ion forming a guanine Hoogsteen-edge contact.

### *MG_diagnosis* (script 1): identifying/correcting/validating/annotating Mg^2+^ ions

All *Cat_Wiz* scripts, developed for *Coot* version 0.9.8.98, generate *GtkTables.* The table produced by the *MG_diagnosis* script provides precise stereochemical assessments of structural Mg^2+^ ions to enable the discrimination between improperly from properly assigned ions (**Figure 4**). The table is organized into multiple tabs that guide users through inspection, validation and refinement. The first two tabs can be exported as a comma-separated value (CSV) file.

- *Tab1 (Mg^2+^ assignment issues) and tab2 (Validated Mg^2+^ ions):* The *“1. Mg^2+^ (check)”* tab identifies ions displaying one or more stereochemical discrepancies. To flag potential issues, three coordination distance ranges are defined: 1.9–2.3 Å, 2.3–2.6 Å and 2.6–3.2 Å. Each range associates with a coordination number (CN) indicator. An ideal Mg^2+^ ion (11,12,14) is characterized by CN = 6 indicating that all six coordination distances fall within the 1.9–2.3 Å range (**Figure 2**). Such ions are listed in the *“2. Mg^2+^ (validated)”* tab (**Figure 4**). A CN = 5/1/0 reflects the presence of a slightly out-of-range coordination distance, while CN = 0/0/5 suggests a non-Mg^2+^ solvent particle, often consistent with K^+^ when present in the experimental buffer (11,14,57). Re-inspection and local refinements are often necessary to validate the assigned ion identity. Color-coding facilitates interpretation: red highlights ions with coordination distances in the 2.6–3.2 Å range typical of K^+^ (tab1), while green marks validated octahedral Mg^2+^ ions with CN = 6 (tab2).
- *Binding site quality:* Each Mg^2+^ binding site is evaluated using three quality metrics *“rmsd ion”*, *“distorted octahedron (Dist. oct.)”* and *“Clash”* which together define the *“high-”, “fair-” and ‘low-quality”* tags. The cells in these columns are color-coded in grey (low), pink (fair) or green (high) to facilitate a rapid assessment.
- *“rmsd” indicator:* The *“rmsd”* value calculated by *Coot* is used as a primary quality marker. Sites with rmsd values exceeding 10% of the highest peak are considered high-quality (*green*). Values below 5% are tagged as *“low-quality”* (*grey*) while intermediate values (5–10%) are deemed *“fair”* (*pink*). Map-dependent rmsd thresholds are noted in tab5 (*“Map info”)*. Importantly, rmsd values are computed from the original experimental map rather than from the smoothed map.
- *Distorted coordination octahedron (Dist. oct.) indicator:* This indicator evaluates distortion of the Mg^2+^ coordination geometry by measuring distances between first-shell atoms which average 2.9 Å in an ideal octahedron (**Figure 2A**). As a proxy for overall coordination quality, only the shortest of these inter-atom distances is reported. Sites are classified as *“high-quality”* when this distance exceeds 2.6 Å (green), *“fair-quality”* between 2.4 and 2.6 Å (pink), and *“low-quality”* below 2.4 Å (grey; **Figure 5**).
- *Clash indicator:* Strict Mg^2+^ coordination distances (< 2.3 Å) generate a 2.3–3.4 Å exclusion zone around the ion (**Figure 2B**). The presence of atoms within the 3.2–3.4 Å range induces a severe clash (grey). In some convoluted binding patterns, distances of 3.4–3.6 Å are categorized as mild clashes (pink). Such clashes generally require deletion or reassigment of the Mg^2+^ ion, remodeling of the binding site, or re-evaluation of ion identity. Representative examples are discussed in the **Results** section.
- *Global quality indicator:* A global quality score is derived by aggregating the individual indicators described above. If any single metric indicates *“low-quality”*, the entire site is classified accordingly. Otherwise, sites are labeled as *“fair-”* or *“high-quality”*. This global indicator applies only to *“validated”* Mg^2+^ ions in tab2 and is reported in the *“Quality”* column. This indicator is not relevant for the non-octahedral coordination patterns noted in tab1.
- *Visualization and remodeling of Mg^2+^ binding sites:* A pop-up window, accessible by a right-mouse click on a tab1 or tab2 line, provides visualization and refinement options (**Figure 4D**). Selecting *“Zoom to this ion”* centers the view on an ion and fascilitates inspection of *d*(Mg^2+^…O/N) distances, missing water molecules, or incomplete octahedral shells. Slightly elongated coordination distances can often be corrected through local real-space refinement in *Coot*, while missing water molecules can be added using density-guided placement tools of *Coot*. When solvent densities are located at more than 4.3 Å from solute atoms and lack clear octahedral density patterns, distinguishing hydrated Mg[H_2_O]_6_^2+^ ion from alternative solvent particles becomes challenging. In such cases, careful examination of second-shell contacts, steric clashes, and experimental buffer composition is required to validate ion identities (14).
- *Spherical refinement of the ion binding site: Coot* real-space refinement tools can be used at any stage. The *“sphere refine”* command, accessible by a right-mouse click on a tab1 or tab2 entry, performs unrestrainted real-space refinement of atoms within specified radius (default = 3.5 Å). When density patterns are well-defined, this typically yields near-ideal octahedral geometries. However, in regions with blurred densities, unrestrained refinement may introduce distortions (**Figure 6**). In such cases, the *“sphere refine with restraint”* option is recommended, as it enforces 2.07/2.19 Å restraints for the six *d*(Mg^2+^…O/N) coordination distances.
- *Additional distance restraints:* In some instances, and for less-well resolved sites, additional restraints may be required to stabilize the binding site and avoid too-close Mg^2+^…O/N contacts. This can be done by navigating to the *“Calculate* → *Scripting* → *Python”* menu using the *Coot* command *“add_extra_bond_restraints”*, which requires thirteen arguments as listed in the **SI**.
- *Mg^2+^…Mg^2+^/K^+^ ion pairs:* When two residues coordinate the same Mg^2+^ ion, the “*sphere refine with restraint”* option may induce slight deviation from ideal 2.07/2.19 Å coordination distances, typically remaining within *d*(Mg^2+^…O/N) < 2.3 Å. Such deviations are commonly observed in Mg^2+^…Mg^2+^ and other ion pairs, which require a dedicated inspection. *MG_diagnosis* identifies all Mg^2+^…Mg^2+^/K^+^ pairs with inter-ion distance ≤ 6.0 Å. Pairs with *d*(Mg^2+^… Mg^2+^/K^+^) < 4.2 Å (orange) are currently poorly documented and require careful analysis, while those in the 4.2–6.0 Å range (cyan) should be visually inspected to assess potential Mg^2+^ μ-clusters involving shared phosphate atoms (14).
- *Doted spheres to visualize clashes:* For ambiguous cases, *MG_diagnosis* allows visualization of doted spheres with radii of 2.3, 3.2, and 3.6 Å. A site is considered stereochemically valid if all six coordinating atoms fall within the 2.3 Å sphere. Atoms within the 2.3–3.2 Å range indicate the need for correction, while solute atoms penetrating the 3.2 Å sphere are incompatible with hydrated Mg^2+^ ion requires that no solute atoms enter the 3.2 Å sphere. Ideally, no atoms should enter the 3.6 Å sphere, as second shell coordination distances peak around this value (**Figure 2B**).
- *Tab3 summary:* The third tab *“3. Mg^2+^ (summary)”* provides an overview of the principal characteristics of validated ions including a list of their binding site types. Together, the first three tabs provide an appropriate overview of the quality of the Mg^2+^ assignments for users who do not intend to further refine the ionic coordination shells.

**Figure 4.**
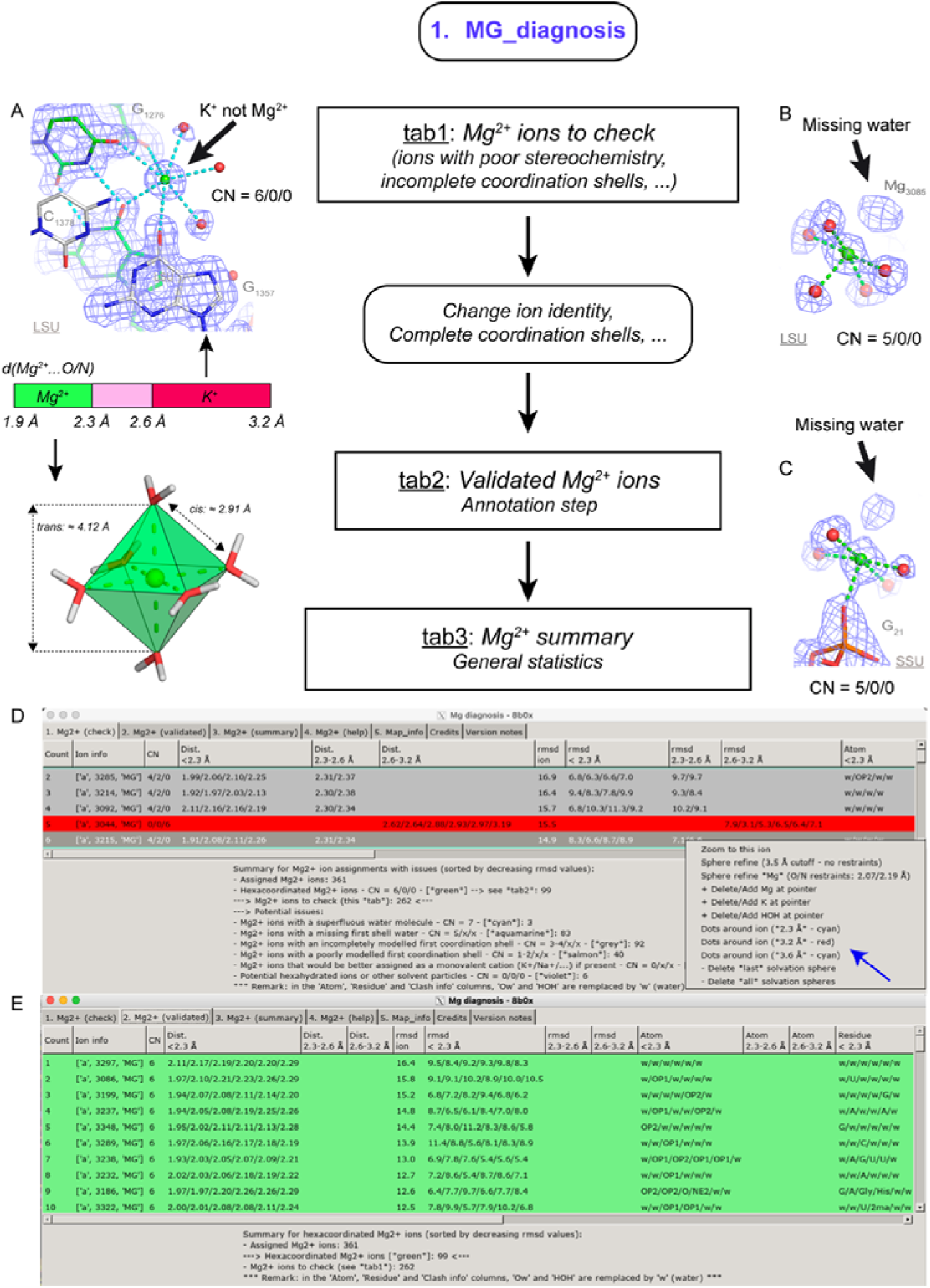
Workflow of the *MG_diagnosis* script. The central column illustrates the *MG_diagnosis* primary functions, i.e., *i)* assessing the stereochemical quality of assigned ions and *ii)* recommending changes based on specific coordination number (CN) and distance criteria. These examples are taken from the **8b0x** structure. (**A**) A color-coded scale is used to evaluate coordination distances. Green corresponds to an ideal 1.9–2.3 Å range for Mg^2+^, pink to an improbable 2.3–2.6 Å range and red to a 2.6–3.2 Å range characteristic of misassigned K^+^ ions. The top-left panel shows a misidentified K^+^ ion with CN = 6/0/0 (14). (**B, C**) Two Mg^2+^ ions with CN = 5/0/0, pointing to a missing first shell water as exposed by the density map. (**D**) Partial view of the *MG_diagnosis* tab1, which lists all identified issues for a given structure. A pop-up menu (blue arrow) accessible through a right-mouse click, provides context-specific options. (**E**) Partial view of the *MG_diagnosis* tab2 (the tab1 and tab2 extensions to the right are not shown).

**Figure 5.**
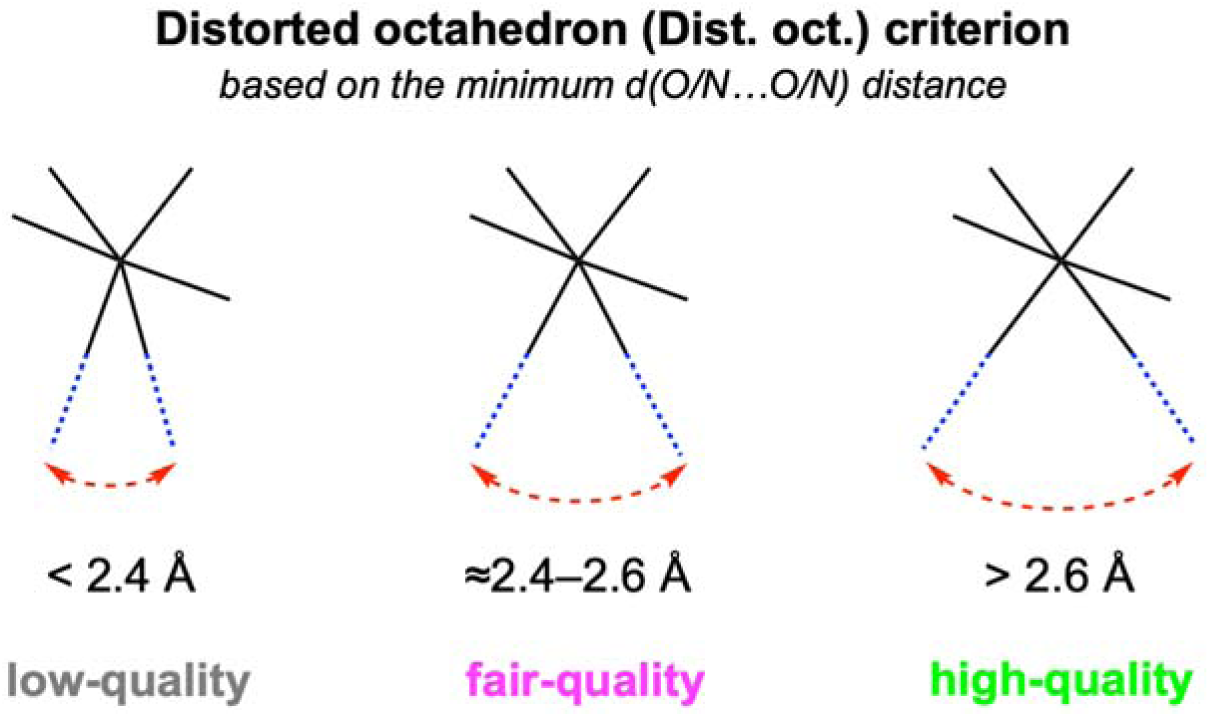
The distorted octahedron (Dist. oct.) criterion. The quality of a binding site is *“low”*, *“fair”,* or *“high”* based on the smallest distance between any two first shell coordinating atoms. This criterion is exclusively applied in tab2 of *MG_diagnosis*. These distance cutoffs were empirically inferred.

**Figure 6.**
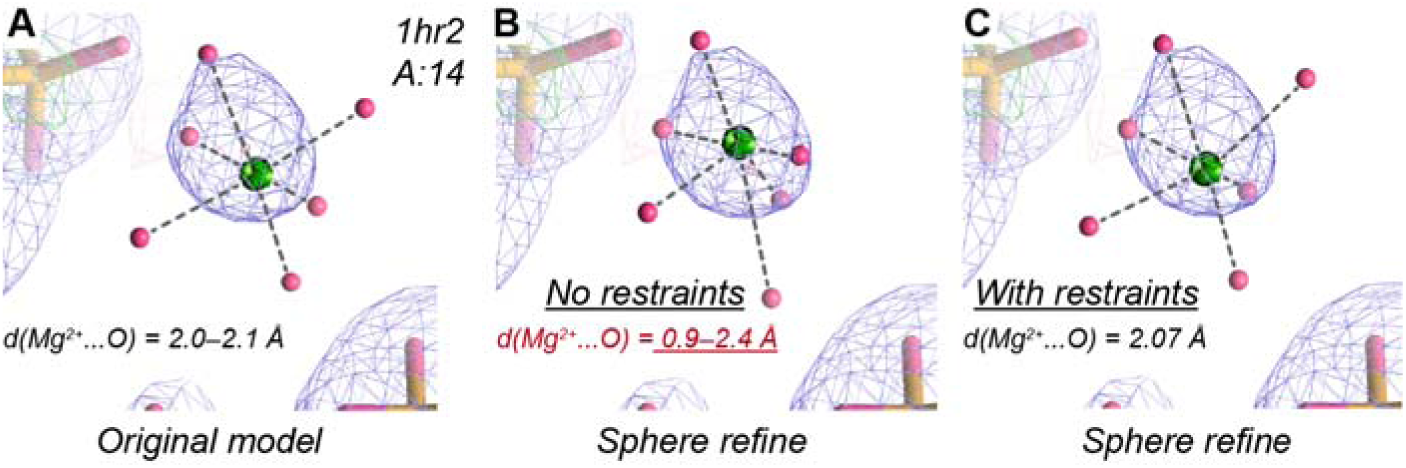
Effect of restraints on the *Coot “Sphere refine”* option. Different outcomes are observed when using the *Coot “Sphere refine”* command with and without distance restraints. (**A**) A hexahydrated Mg^2+^ ion modelled into a non-octahedral density pattern (89). (**B**) Refining the same position using the *“sphere_refine”* command without added restraints shows a deformed coordination octahedron with out-of-range coordination distances. (**C**) When the refinement is performed with added restraints, the octahedral geometry of the hydrated Mg^2+^ ion is preserved, despite minor deformation.

*Tab4 (Help) and tab5 (Map info):* The fourth and fifth tabs, *“4. Mg^2+^ (help)”* and *“5. Map info”,* provide general information on the *MG_diagnosis* scripts and the experimental maps, respectively.

### *MG_detect* (script 2): localizing unassigned Mg^2+^ ions

While *MG_diagnosis* annotates all assigned Mg^2+^ ions, it is not designed to locate unassigned ions. This task is devoted to the *MG_detect* and *MG_clamp* modules which are ideally run in the order shown in **Figure 1**. *MG_detect* comprises *MG_detect_RNA* and *MG_detect_Prot*. These scripts use the *Coot “mask_map_by_molecule”* command with an atom radius of 1.6 Å to create a masked map around the selected molecule and thereby isolate unassigned density spots (**Figure 7**).

**Figure 7.**
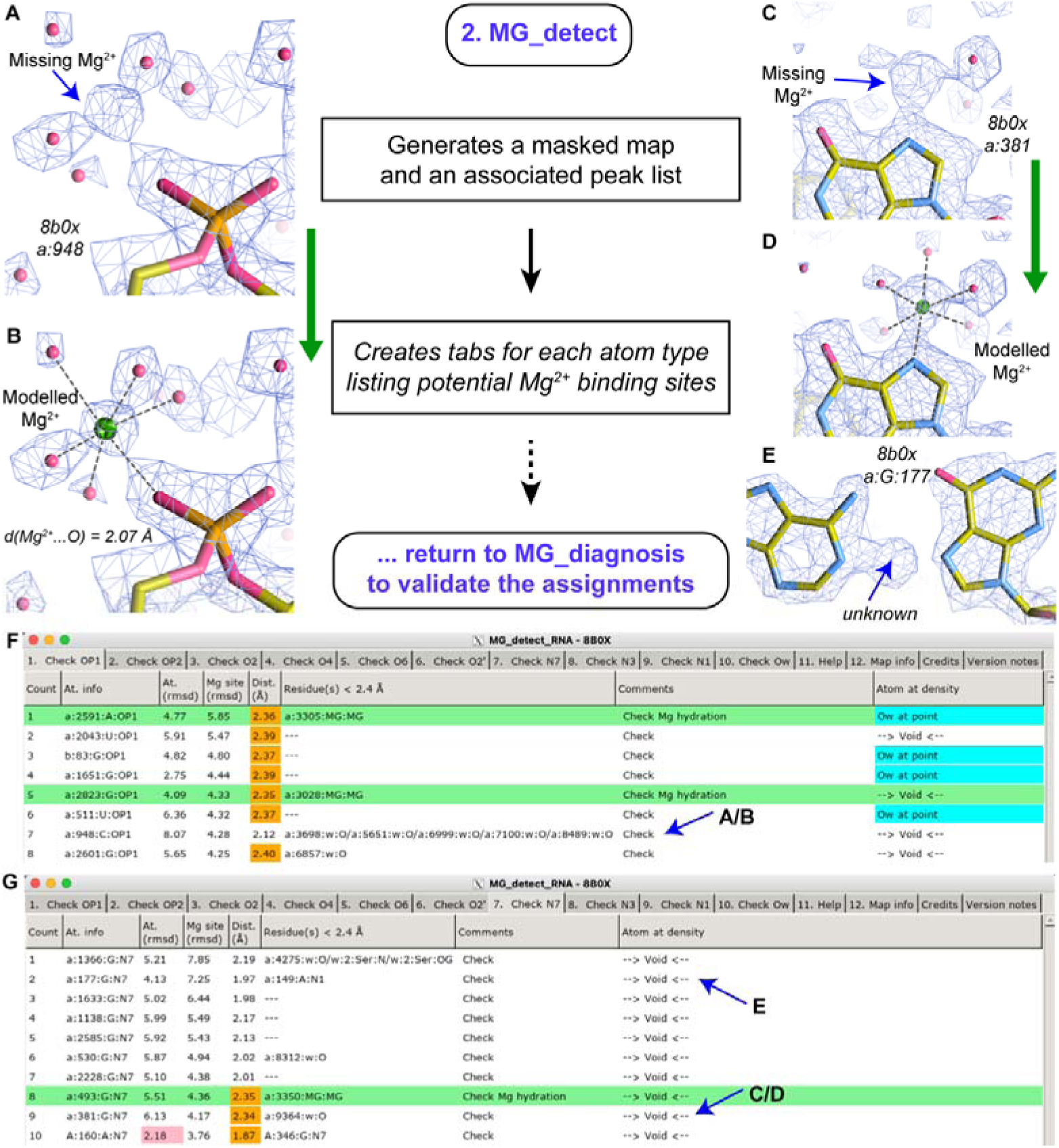
Organization of the *MG_detect_RNA* script. *MG_detect_RNA* identifies potential Mg^2+^ binding sites by collecting the highest non-assigned peaks from a masked map. (**A**) Shows a missing Mg^2+^ ion that is part of an O_ph_.5O_w_ binding site successfully identified by *MG_detect_RNA*. (**B**) Shows subsequent modelling of this site using *MG_diagnosis* options. (**C/D**) Show a similar process for a missing Mg^2+^ that is part of an N_7_.5O_w_ site. (**E**) Shows density associated with an unidentified ion that is part of a metal-mediated A…G pair as described in (14). (**F/G**) Show partial views of the generated tabs which identify potential Mg^2+^ binding sites near OP1 and N7 atoms, respectively (marked by blue arrows). Orange cells mark distances from the identified density peak to the selected atom that are in the 2.3–2.4 Å range or below 1.9 Å; pink cells mark low rmsd values.

*MG_detect_RNA* and *MG_detect_Prot* inspect density spots in proximity to nucleotide and protein atoms. For nucleotides, these include phosphate and nucleobase atoms (OP1/OP2/O2/O4/O6/O2’/N7/N3/N1), while for proteins they include side-chain atoms (OD1/OD2/OE1/OE2/ND1/NE2/N/NZ/NE/NH1/NH2) as well as backbone atoms (O_bb_). Note that the nucleobase N1/N3 and protein N/NZ/NE/NH1/NH2 atoms are not involved in Mg^2+^ binding but can occasionally coordinate transition metals as detailed below. For modified nucleotides or amino acids that involve different atom names, nearby density spots must be inspected manually. Since *MG_detect_RNA* and *MG_detect_Prot* operate analogously, the description below focuses on *MG_detect_RNA*.

The generated mask is searched for the highest-density peaks using *Coot “map_peaks_around_molecule”* command. In all tabs associated with OP1/OP2/O2/O4/O6/O2’/N7/N3/N1 atoms, peaks are ranked by decreasing rmsd values in the *“Mg site”* column. To avoid overlaps with existing atoms, only peaks located within 1.8–2.4 Å of any assigned atom are retained as candidates for inner-sphere contacts. The *“Residues < 2.4 Å”* column lists all residues located near each potential *“Mg site”* peaks. Thus, peaks within 1.8–2.4 Å of an O/N atom are listed as potential inner-sphere Mg^2+^ ion locations.

The lower-ranked peaks (rmsd below 5% of the highest peak value) inevitably include spurious positions caused by map irregularities or noise rather than genuine solvent particles. The number of these peaks depends on the treatment (smoothing or sharpening) applied to the map (58,59). Thus, peaks with rmsd values below 5% of that of the highest peak are excluded while peaks in the 5–10% range are shown in pink.

The highest-ranked peaks (rmsd above 10% of the highest peak value) are typically associated with unassigned Mg^2+^ ions that directly contact RNA atoms (**Figures 7A–D**). However, some peaks may correspond to *“unknown”* atoms, such as those previously identified in **8b0x** that appear to be metal-mediated base pair components (14). For instance, in **8b0x**, the top-ranked peak associated with N7 atoms cannot be associated with Mg^2+^, K^+^ or water and instead indicates a metal-mediated base pair interaction (**Figures 7E–G**). Similarly, in **9q87**, we tentatively used the “CU” PDB residue identifier that corresponds to copper, based on the rationale that copper grids were employed and that these ions typically exhibit mostly in-plane coordination with CN = 4. Peaks near nucleobase N3/N1 atoms do not indicate Mg^2+^ binding sites but may reveal the presence of transition metals or problematic local modeling.

Interestingly, some peaks also highlight poorly modelled backbones or reveal the presence of alternate conformations. We therefore intentionally avoid excessive filtering this list to allow such issues to be detected and corrected. In addition, *MG_detect* can also locate unassigned densities consistent with octahedral or tetrahedral transition metals (Mn^2+^, Zn^2+^, etc.) that share Mg^2+^ coordination distances. As a note of caution, these modules also report peaks that associated with diffuse or ambiguous densities, which may arise from disorder, unidentified solvent particles or insufficient knowledge.

Besides inner-shell contacts, *MG_detect_RNA* can identify hexahydrated Mg^2+^ ions of the 6O_w_ type, provided that water molecules have been assigned. For that, automated procedures implemented in *Coot* or *Phenix* are recommended. Different cutoffs might be used for an optimal screening. In these instances, *MG_detect* locates densities close to already assigned water molecules with distances below 2.4 Å. The list of peaks is provided in the *“Check O_w_”* column.

In summary, *MG_detect_RNA* and *MG_detect_Prot* identify unassigned density peaks that may correspond to Mg^2+^ ions or transition metals while excluding peaks associated with water and monovalent ions (e.g., K^+^ or NH_4_^+^) that exhibit coordination distances in the 2.6–3.2 Å range. Following *MG_detect,* all newly assigned Mg^2+^ positions must be evaluated using *MG_diagnosis* to validate ion identity, correct stereochemistry and complete their coordination shell.

### *MG_clamp* (script 3): detecting and annotating Mg^2+^ clamps

Several techniques are available to detect Mg^2+^ binding clamps (**Figure 2**). For instance, *MG_detect_RNA* captures densities near O_ph_/O_b_/O_r_/N_b_ atoms that can subsequently be assigned to Mg^2+^ ions. This approach is highly effective for most high-resolution structures (< 2.0 Å). However, in lower-resolution structures or in poorly resolved regions of large complexes, the experimental density patterns may be insufficiently defined for *MG_detect* to be fully reliable. In such cases, *MG_clamp provides* a complementary identification strategy by searching for pairs of atoms with *d*(O/N…O/N) ≤ 3.4 Å (**Figure 8**). This helps locating Mg^2+^ binding motifs with weak ion densities but sometimes also identifies poorly modelled RNA backbones. Worth noting, two non-protonated O/N atoms separated by *d*(O/N…O/N) ≤ 3.4 Å can only be stabilized by coordination to Mg^2+^ or to other transition metals. Such stabilization typically occurs via two direct O/N…Mg^2+^ contacts or more rarely through one direct and one water-mediated contact (**Figure 3**). To date, no examples of clamp motifs stabilized by monovalent ions have been reported (8,30).

- *Tab1 (“Fix Asn/Gln” side chains):* Misoriented protein side chains can occasionally generate artefactual contacts that mimic genuine Mg^2+^ clamps with *d*(O…O) ≤ 3.4 Å. Even after refinement using appropriate protocols in *Phenix* or *Molprobity* (60,61), Asn or Gln side chains may revert to an inappropriate orientation (**Figures 8A/B/E**). *MG_clamp* therefore identifies all *d*(O/N…O/N) ≤ 3.4 Å distances involving Asn/Gln side chains. When a misorientation is detected, it is recommended to invert the Asn/Gln(N/O) positions with the *Coot “flip 180°”* tool to improve the model.
- *Tab2 (“Potential clamps with no Mg^2+^ density”):* The second tab lists all non-water contacts with d(O/N…O/N) ≤ 3.4 Å that are not yet associated with an Mg^2+^ ion. Because most hydrogen bond contacts fall into this category, several criteria are used to reduce false positive. For example, the identification process prioritizes atom pairs that include at least one O_ph_ atom since over 99% of the clamps detected in **8b0x** belong to this type (**Table 1** exceptions include 2N_b_.4O_w_ and 2O_b_.4O_w_ binding types). Atom pairs with low *rmsd* values are also flagged, as they frequently indicate incompletely modelled sites. These pairs are retained for visual inspection.
- *Tab3 (“Mg^2+^ clamps”):* The third tab identifies and annotates all d(O/N…O/N) ≤ 3.4 Å contacts that are associated with an assigned Mg^2+^ ion. Clamps involving a hexacoordinated Mg^2+^ ion (CN = 6) are highlighted in green, whereas sites requiring adjustments or corrections are shown in cyan. The latter can typically be remodelled straightforwardly in *Coot.* The *“n…n+1”* column identifies clamps in which Mg^2+^ binds to consecutive residues. For some nucleic acids, water-mediated O_ph_.5O_w_ and O_b_.5O_w_ clamps are also included and annotated in the *“Comment”* column (**Figure 3**). Finally, as with all newly assigned ions, visual inspection of the binding site and the corresponding experimental density using *MG_diagnosis* is required prior to validation.
- *Tab4 (Help) and tab5 (Map info):* The fourth and fifth tabs, *“4. Mg^2+^ (help)”* and *“5. Map_info”*, provide general information on the various processes associated with the *MG_clamp* scripts and the experimental maps.

**Figure 8.**
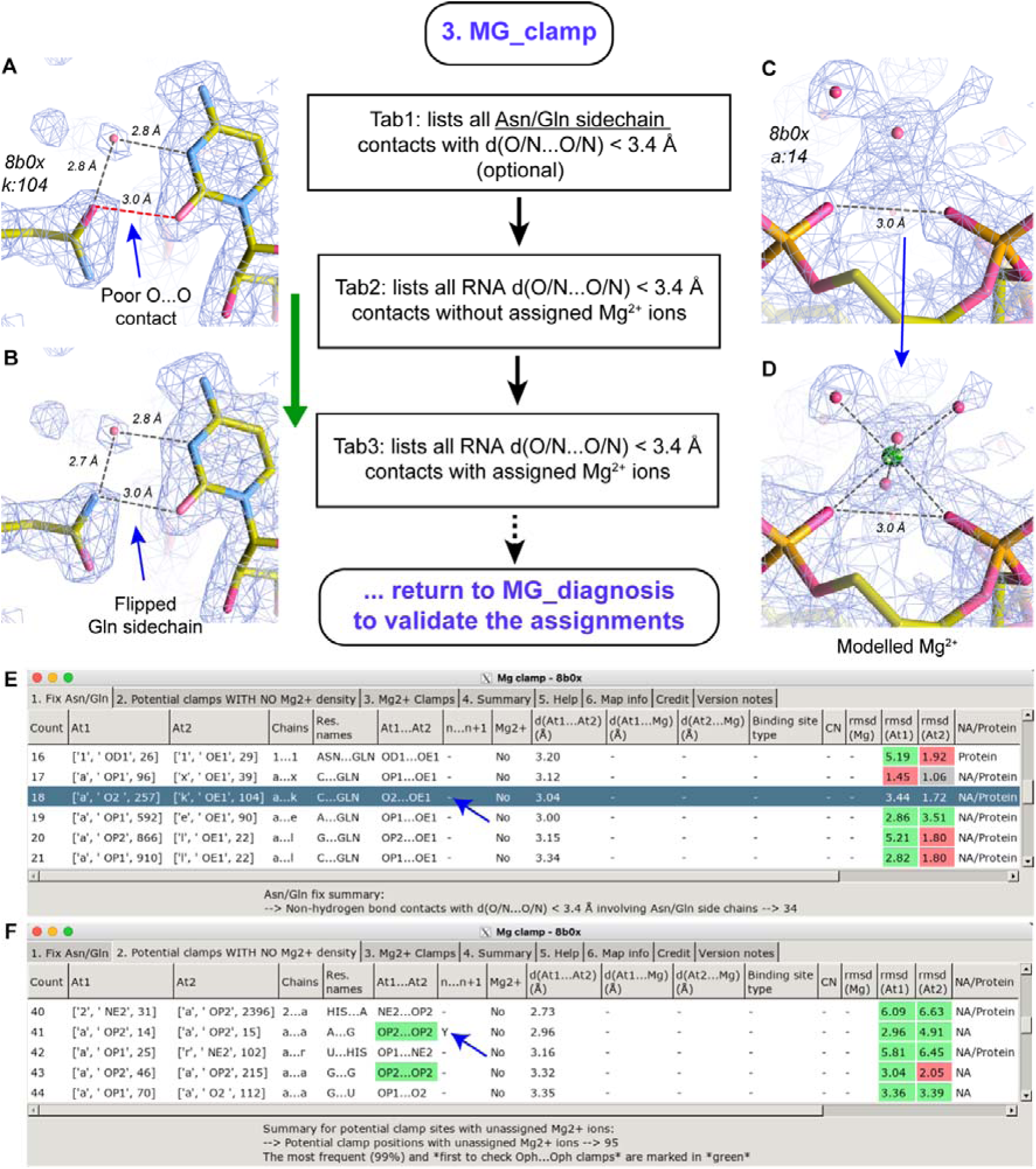
Organization of the *MG_clamp* script. *MG_clamp* locates non-hydrogen-bonded pairs of atoms with *d*(O/N…O/N) below 3.4 Å. These pairs are common to all higher-binding motifs (except *trans-*2O_ph_.4O_w_ and related; see Figure 3). When such pairs are identified, a visual inspection of the associated density patterns is required to determine whether the site can accommodate a hydrated Mg^2+^ ion. (**A**) Shows a misoriented Gln side chain that creates an unrealistic *d*(O…O) contact near 3.0 Å. (**B**) Shows the correct side chain orientation obtained by using the *Coot “flip_sidechain_180°”* tool. (**C**) Illustrates how a 2O_ph_.4O_w_ binding pattern with *d*(O…O) ≈ 3.0 Å can be identified by *MG_clamp*. (**D**) Shows the modelled binding pattern. (**E/F**) Show a partial view of “tab1” and “tab2” generated by *MG_clamp*. In panel **E**, a blue arrow marks the short *d*(O…O) distance caused by an incorrectly oriented Gln side chain (see **A/B**). In panel **F**, a blue arrow marks a short *d*(O…O) distance detected by *MG_clamp* that is associated with an unassigned *cis-*2O_ph_.4O_w_ Mg^2+^ ion (see **C/D**).

**Table 1.** Occurrence of well-defined RNA octahedral Mg^2+^ binding sites in the 8b0x and 9q87 structures. *“High”, “fair”* and “*low”* quality binding site occurrences are provided in parenthesis.

|  | 8b0x |  | 9q87 |  |
| --- | --- | --- | --- | --- |
| **Outer-shell** |  |  |  |  |
| 6O<sub>w</sub> | 21 (17/1/4) |  | 118 (48/53/17) |  |
| **Pentahydrated** |  |  |  |  |
| O<sub>ph</sub>.5O<sub>w</sub> | 38 (28/7/3) |  | 149 (71/57/21) |  |
| O<sub>b</sub>.5O<sub>w</sub> | 3 (3/–/–) |  | 16 (8/7/1) |  |
| O<sub>r</sub>.5O<sub>w</sub> | 1 (1/–/–) |  | 1 (–/1/–) |  |
| N<sub>b</sub>.5O<sub>w</sub> | – |  | 9 (1/5/3) |  |
| O<sub>coo</sub>.5O<sub>w</sub> (with r-protein) | – |  | 1 (1/–/–) |  |
| **Tetrahydrated** | *cis*- | *trans*- | *cis*- | *trans*- |
| 2O<sub>ph</sub>.4O<sub>w</sub> | 12 (9/2/1) | 3 (2/–/1) | 46 (25/12/9) | 15 (4/6/5) |
| 2O<sub>b</sub>.4O<sub>w</sub> | – | – | 1 (–/1/–) | – |
| 2N<sub>b</sub>.4O<sub>w</sub> | – | – | 3 (1/2/–) | – |
| O<sub>ph</sub>.O<sub>b</sub>.4O<sub>w</sub> | 3 (2/–/1) | 1 (–/–/1) | 10 (7/1/2) | 1 (1/–/–) |
| O<sub>ph</sub>.N<sub>b</sub>.4O<sub>w</sub> | – | – | 2 (2/–/–) | 2 (1/–/1) |
| O<sub>ph</sub>.N<sub>his</sub>.4O<sub>w</sub> (with r-protein) | 1 (1/–/–) | – | 2 (2/–/–) | – |
| **Trihydrated** | *fac*- | *mer*- | *fac*- | *mer*- |
| 3O<sub>ph</sub>.3O<sub>w</sub> | 3 (2/1/–) | 6 (6/–/–) | 7 (5/2/–) | 9 (9/–/–) |
| 2O<sub>ph</sub>.O<sub>b</sub>.3O<sub>w</sub> | – | 1 (1/–/–) | – | 2 (2/–/–) |
| 2O<sub>ph</sub>.O<sub>r</sub>.3O<sub>w</sub> | – | – | 1 (1/–/–) | – |
| 2O<sub>ph</sub>.N<sub>b</sub>.3O<sub>w</sub> | 1 (1/–/–) | – | 2 (2/–/–) | – |
| O<sub>ph</sub>.2O<sub>b</sub>.3O<sub>w</sub> | – | – | – | 1 (–/1/–) |
| 2O<sub>ph</sub>.O<sub>cno</sub>.3O<sub>w</sub> (with r-protein) | – | 1 (1/–/–) | – | 1 (1/–/–) |
| **Dihydrated** | *CIS*- | *TRANS*- | *CIS*- | *TRANS*- |
| 4O<sub>ph</sub>.2O<sub>w</sub> | 1 (1/–/–) | 1 (1/–/–) | 1 (1/–/–) | 1 (1/–/–) |
| 3O<sub>ph</sub>.O<sub>b</sub>.2O<sub>w</sub> | – | – | 1 (1/–/–) | – |
| 3O<sub>ph</sub>.O<sub>r</sub>.2O<sub>w</sub> | – | – | 1 (1/–/–) | – |
| 2O<sub>ph</sub>.2O<sub>b</sub>.2O<sub>w</sub> | 1 (–/–/1) | – | 1 (–/1/–) | – |
| 2O<sub>ph</sub>.O<sub>bb</sub>.N<sub>his</sub>.2O<sub>w</sub> (with r-protein) | 1 (1/–/–) | – | 1 (1/–/–) | – |
| O<sub>ph</sub>.3O<sub>bb</sub>.2O<sub>w</sub> (with r-protein) | – | – | 1 (–/–/1) | – |
| 3O<sub>coo</sub>.N<sub>his</sub>.2O<sub>w</sub> (r-protein) | – | – | 1 (–/1/–) | – |
| **Sub-total:** | 99 (76/12/11) |  | 406 (197/150/60) |  |
| **With issues** (not in table): | 262 |  | 9 |  |
| **Total:** | 361 |  | 415 |  |

### Updating B-factors at the end of the assignment process

If new Mg^2+^ assignments are introduced or existing ion positions are modified, *B-factors* must be updated prior to PDB deposition. This can be achieved by refining individual *B-factors* using *Phenix* or an equivalent refinement program. Care is required when working with over-sharpened maps, which may otherwise lead to zero or negative *B-factors* during refinement.

## RESULTS

### Case I: The 1.55 Å *Escherichia coli* ribosome

Analysis of the *E. coli* ribosome structure determined at 1.55 Å resolution (PDBid: **8b0x**) serves as the primary case study for evaluating the performance *Cat_Wiz*. This structure was previously investigated in detail by us, revealing the presence of 361 Mg^2+^ ions (14,19). A comprehensive re-inspection of the structure led to the reassignment and validation of 403 Mg^2+^ ions, a process that formed the basis for establishing a systematic classification of Mg^2+^ binding sites in RNA and RNA/protein systems (14). Besides identifying new binding sites, this earlier study corrected misassignments by reclassifying several Mg^2+^ ions as water molecules or K^+^ ions and by deleting others. Using *Cat_Wiz*, we re-examined the previously **revised_8b0x** structure. This second revision identified seven additional Mg^2+^ binding sites, while validating the positions of existing Mg^2+^ ions in the amended structure in just a few minutes – a task that otherwise requires days of manual inspection. No novel Mg^2+^ binding motifs beyond those summarized in **Figure 3** were detected during this process. The revised structure, deposited in the PDB under the identifier **9q87**, reveals significant differences in the ionic shell compared with the original **8b0x** and the new structure (**Table 1**). In **8b0x,** only 100 Mg^2+^ ions exhibit a hexacoordinated geometry that met our stereochemical criteria. In contrast, the refined **9q87** structure contains 406 validated hexacoordinated Mg^2+^ binding sites.

*MG_diagnosis* proved essential for guiding this revision process. The script flagged 261 ions requiring re-evaluation of their chemical identity or their coordination shell. Of those, 42 ions were identified as potentially misassigned K^+^ ions or other solvent particles, while 220 ions required coordination-shell adjustments. Among the latter group, 85 sites needed only minor corrections (e.g., coordination distance adjustments), whereas the remaining 135 sites required a complete remodeling. The ability of *Cat_Wiz* to diagnose such issues rapidly and to facilitate their correction significantly streamlined the refinement process, even for users with limited prior experience with *Coot*.

MG_detect further improved completeness of the **amended 8b0x** (14) model by identifying 12 additional Mg^2+^ ions while nine previously assigned Mg^2+^ ions were flagged for anomalous features (**Table 1**). One puzzling ion exhibited a coordination shell of seven water molecules with coordination distances close to 2.07 Å, a geometry more consistent with a K^+^ assignment than Mg^2+^. Eight additional ions were associated with Mg^2+^…Mg^2+^ ion pairs and displayed atypical coordination patterns. Approximately 50 *“low-quality”* instances were identified (**Table 1**). Although these sites passed inspection and did not exhibit steric clashes, they are not suitable for inclusion in statistical surveys or AI training sets. Nevertheless, documenting these sites remains important for achieving a complete description of the ribosome solvent shell. Finally, *MG_detect identified* one ion with a coordination number of five that appears to be directly linked to an alternate conformation of a guanine nucleobase. As illustrated in **Figure 10**, the high-resolution structure reveals two alternate conformations. These conformations are challenging to interpret and remain outside the scope of this study.

We also used *MG_clamp* to identify short *d*(O…O/N) distances (≤ 3.4 Å) involving Asn/Gln residues, as annotated in tab1 (**Figures 8A/B**). This script flagged 33 contacts in **8b0x** that resulted in the flip of at least 16 Asn/Gln side chains. In one instance, we identified a rare 2O_ph_.O_coo_.3O_w_ RNA/protein-Mg^2+^ binding motif involving the small ribosomal subunit, ribosomal protein (r-protein) S7, and a Mg^2+^ ion at position a:3219. MG_clamp required approximately 10 minutes to run on a standard modern laptop.

A few side notes are necessary for a better understanding of the **8b0x/9q87** structures. First, the *E. coli* ribosome sequence used in this structure originates from strain B rather than the more frequently studied MRE 600 (62,63) or K12 strains (64), resulting in systematic residue numbering shifts (14). Second, four nucleobases (A:U:677; A:G:741; A:U:1086; a:U:2316) are anionic (14,65) and cannot currently be annotated using standard PDB residue nomenclature that does not include such “details”. Third, a recurrent a:2050:TKW residue (5-hydroxycytidine) has been annotated for the first time in bacterial ribosome structures (66). Fourth, we tentatively included six CU residues that account for Cu^+^ (PDBid: CU1) or Cu^2+^ (PDBid: CU) at positions (A:A_160_:N7…A:G_346_:N7; a:A_149_:N1…a:G_177_:N7; a:U_1309_:N7…a:C_1608_:N3; a:U_1854_:N3…a:A_1893_:N7) which likely represent metal-mediated base pairs, as well as between a:G_1366_:N7 and the N-terminal w:Ser_2_ of r-protein bl28. An additional CU site was assigned to a density located between v:His_57_:ND1…v:Cys_53_:SG of r-protein L27 (**Fig. S1**). The observed coordination geometries are consistent with Cu^+^ ions(67–70). If present, these Cu^+^/Cu^2+^ ions represent buffer contaminants potentially originating from grid-related contamination. Importantly, *MG_detect* successfully localized these non-Mg^2+^ densities.

Finally, although *Cat_Wiz* identified the majority of prominent Mg^2+^ clamps in this ribosome (14), it remains difficult to recover the location of all 6O_w_ and 5O_w_ sites. Diffuse ion-binding events, remain currently beyond the reach of *Cat_Wiz* and of available experimental approaches.

### Case II: *Cat Wiz* as an independent tool to investigate metal ion chemistry of hammerhead ribozyme

As a second example, *Cat_Wiz* was applied to analyze Mg^2+^ binding sites in hammerhead ribozyme structures deposited to the PDB and listed in **Table 2/S1**. This RNA has long been regarded as the first RNA metalloenzyme. While the earliest published hammerhead structure (PDBid: **1hmh**) did not mention any Mg^2+^ ions (71), a subsequent X-ray structure (PDBid: **1mme**) reported the position of five Mg^2+^ ions supported by biochemical inferences (72) and subsequent molecular dynamics (MD) studies that proposed catalytic roles for these Mg^2+^ ions (76–80). However, later research challenged these early findings, arguing that Mg^2+^ ions are not directly involved in the catalytic mechanism (73–75). To date, structural studies including X-ray crystallography have not provided sufficient evidence for conserved or catalytically essential Mg^2+^ binding sitesin the hammerhead ribozyme. Current hypothesis instead propose an acid-base catalytic mechanism, in which a hydroxyl group within the active site functions as a general acid, likely activated by a nearby Mg^2+^ ion, while a guanine N1 acts as a base (75,81). Consequently, the hammerhead ribozyme is now more commonly classified as an acid-base catalysed ribozyme rather than a true metalloenzyme.

**Table 2.**
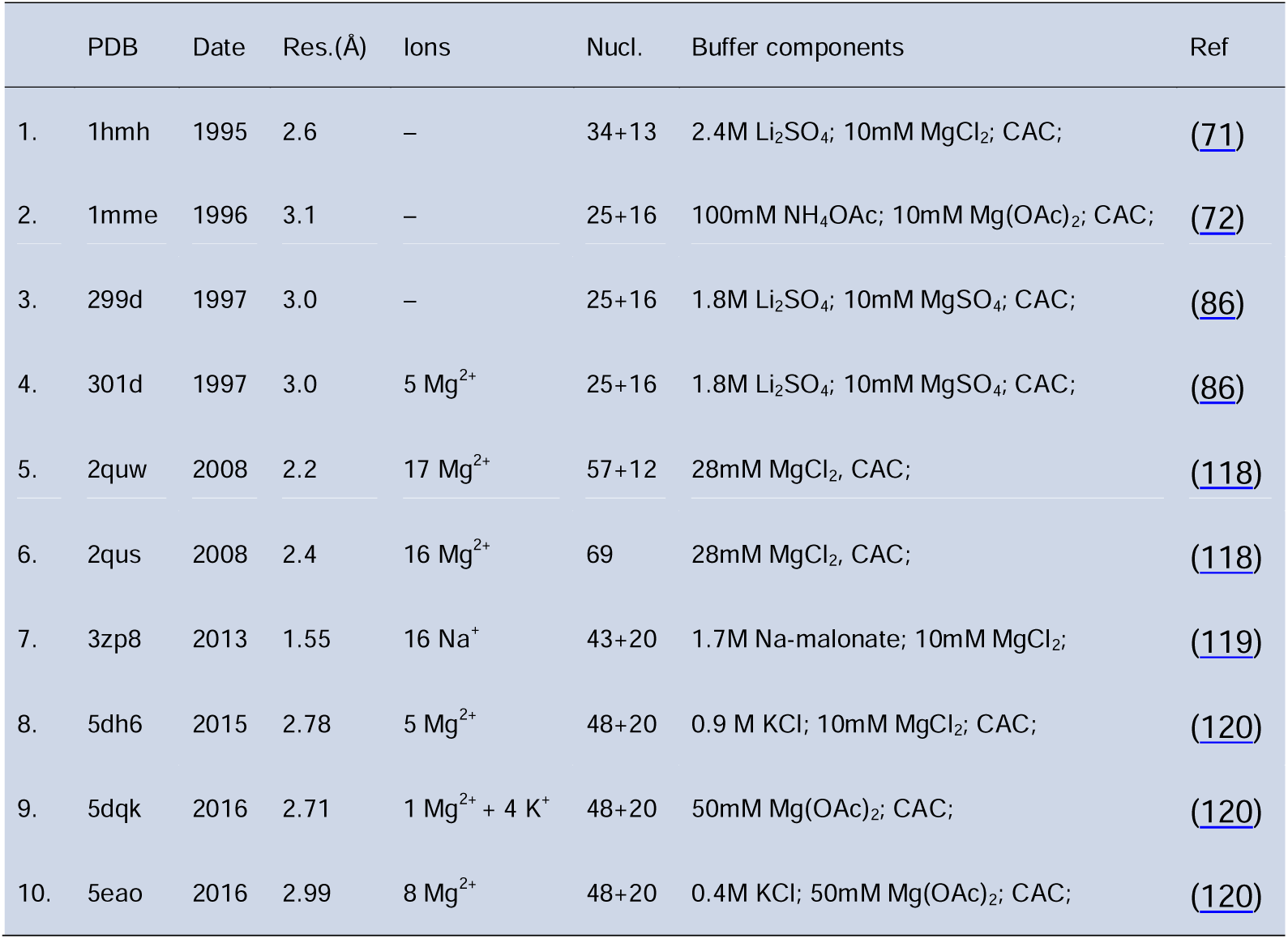
List of X-ray structures of the hammerhead ribozyme with assigned Mg^2+^ ions. The table lists the six PDB structures with assigned Mg^2+^ ions. The associated buffers contain cacodylate or CAC, AsO_2_(CH ) ^−^ anions. Structures with no assigned divalent ions, despite the presence of Mg^2+^ in the buffer, are shown in grey. A complete list of all 26 hammerhead structures is provided in **Table S1**.

A challenge in analyzing these X-ray structures arises from the presence of submolar concentrations of cacodylate (AsO_2_(CH_3_)_2_^−^) and, in some instances, high concentrations of sulfate (SO_4_^2-^) in crystallization buffers (**Tables 2/S1 and Figure 9**). Indeed, several originally assigned Mg^2+^ ions are stereochemically inconsistent with Mg^2+^ coordination and are more plausibly explained as sulfate or cacodylate anions (82,83) which could, in principe, have been detected through their strong anomalous X-ray diffraction signals (84,85).

**Figure 9.**
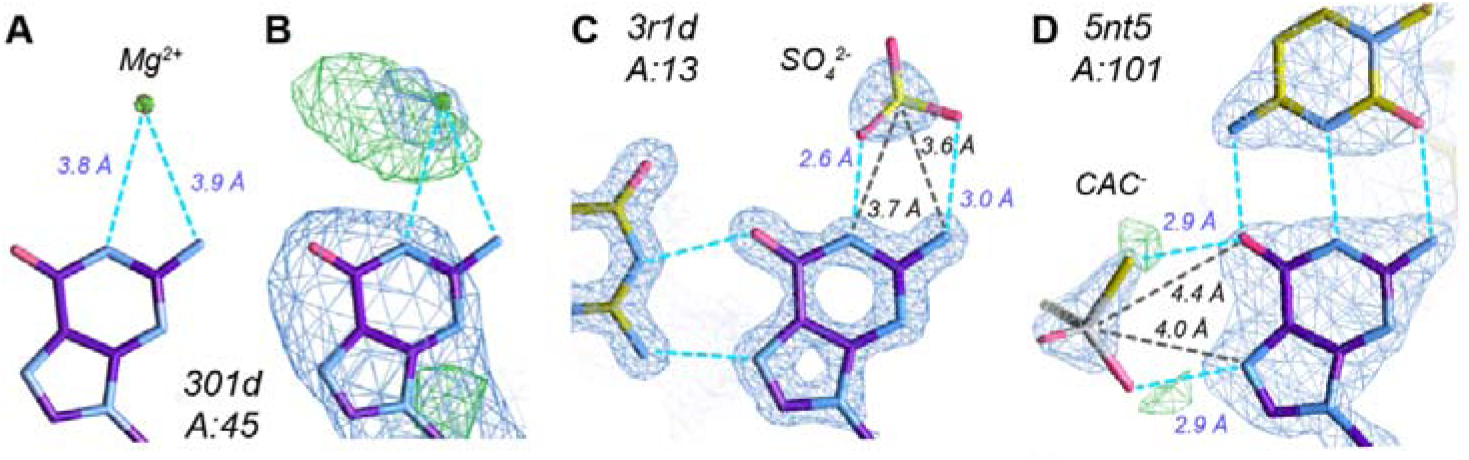
Anionic alternatives to questionable Mg^2+^ assignments. This figure presents examples of alternative explanations to questionable Mg^2+^ assignments. (**A/B**) In the **301d** hammerhead structure, the Mg**^2+^**ion (Mg:45) is unconvincingly positioned near an electropositive Watson-Crick edge (82,83). A more plausible assignment for this density would be a sulfate anion (SO_4_^2-^), as observed in the 1.45 Å resolution structure **3r1d** (**C**), or a cacodylate anion (AsO_2_(CH_3_)_2_^−^), as modelled in the 2.30 Å resolution structure **5nt5** (**D**). Panel **B** shows *fo-fc* density patterns (3.0 σ; green) suggesting the presence of a more electron-dense particle than Mg^2+^. The *2fo-fc* density patterns (1.5 σ; blue) are shown in all panels.

**Figure 10.**
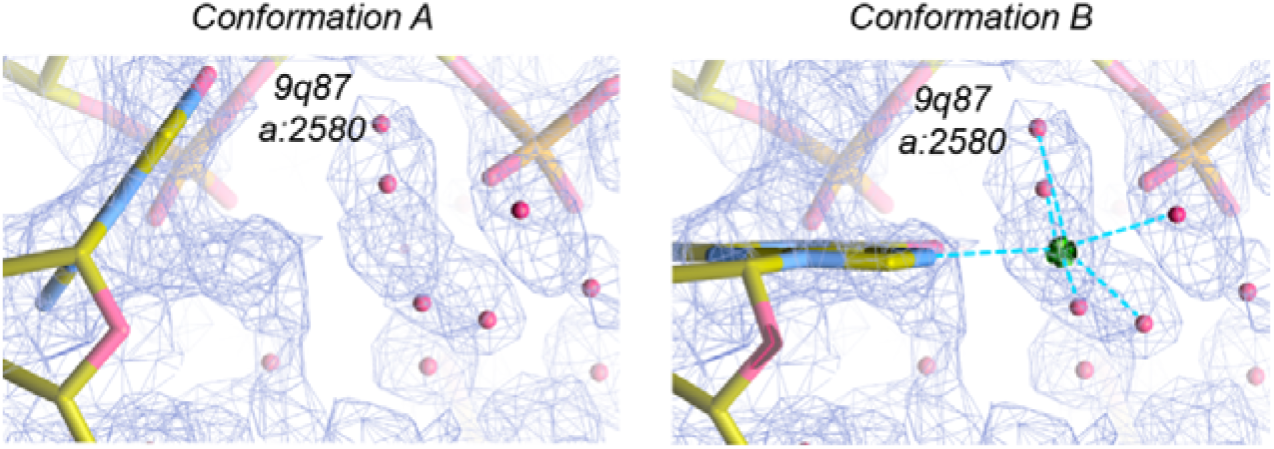
Alternative conformation of the a:2580 residue in 9q87. Conformation A shows the favoured orientation of a:G:2580. Conformation B shows an alternate conformation involving a well-defined N7.5O_w_ contact with Mg^2+^.

To reassess these assignments, we analyzed all structures mentioned in **Table 2** using *Cat_Wiz*. The **1hmh, 1mme,** and **299d** structures lack assigned Mg^2+^ ions although related papers discussed ion dependent catalytic mechanisms (71,72,86). **301d** is the first deposited structure with explicitly assigned Mg^2+^ ions, reporting five such ions. Four of these ions are located more than 3.2 Å away from any nucleotide, consistent with fully solvated ions like Mg[H_2_O]_6_^2+^, although their corresponding densities could equally be attributed to sulfate or cacodylate ions (**Figure 9**). A fifth ion, positioned at a distance of 2.4 Å from A:11:OP2 (**Figure 11A**), is barely supported by experimental density. Interestingly, the assigned occupancy for all five ions is 0.5 while in the parent **359d** structure, Tb^3+^ occupancies are below 0.5 indicating unresolved refinement issues. For all those structures, *Cat Wiz* was unable to propose any strong Mg^2+^ binding site (75,81).

**Figure 11.**
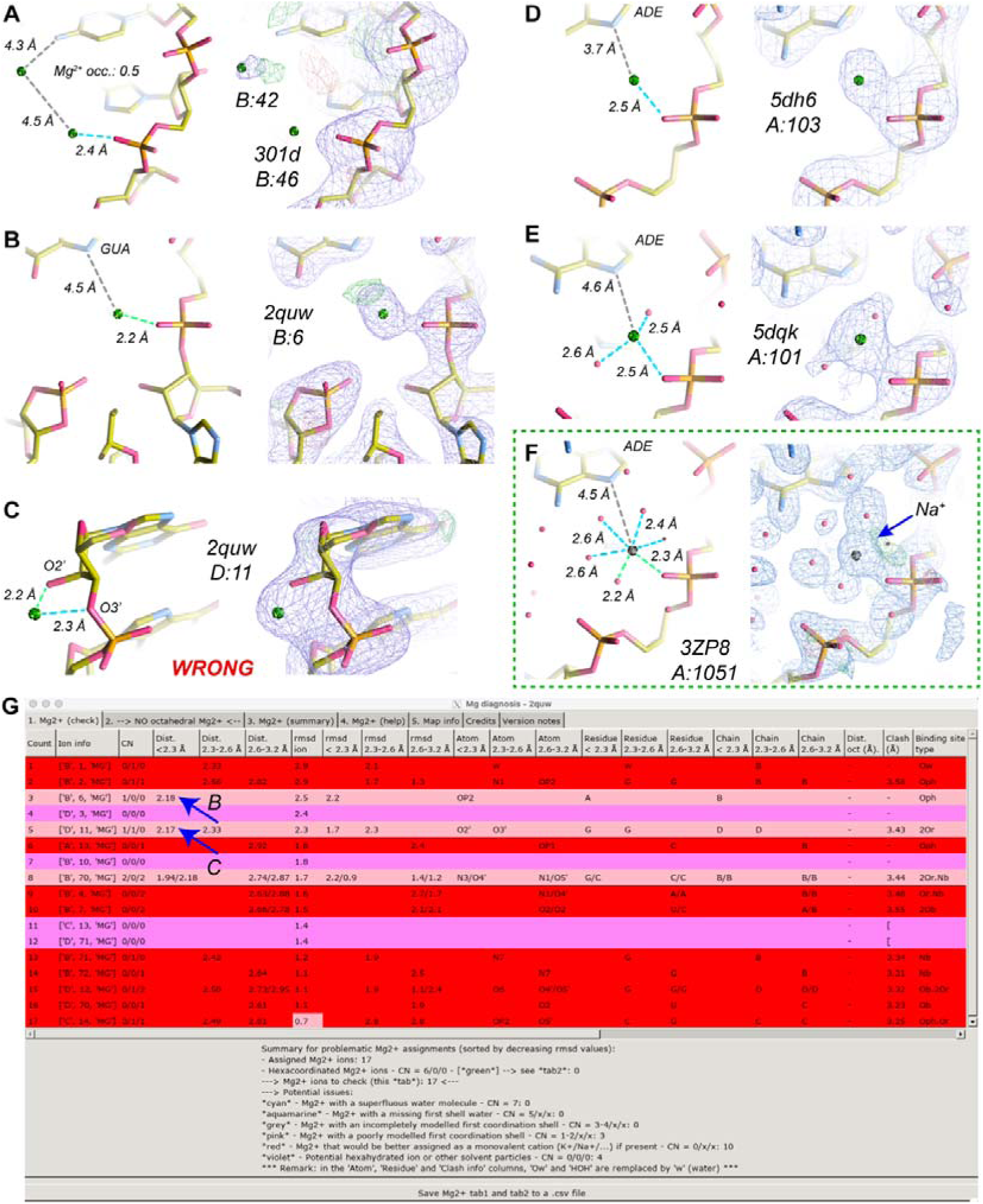
Ion binding assignments in hammerhead ribozyme X-ray structures. (**A**) Mg^2+^ ion tentatively assigned to a weak density spot in the 3.0 Å resolution **301d** structure (occupancy = 0.5). (**B**) A more realistic Mg^2+^ ion from the 2.2 Å resolution **2quw** structure, although lacking octahedral density. (**C**) A stereochemically improbable Mg^2+^ ion. (**D,E**) Two similar assignments from the 2.78 and 2.71 Å resolution **5hd6** and **5dqk** structures, respectively, neither of which display octahedral densities. In **5dqk**, the *d*(O_ph_…O_w_) > 2.6 Å distances further cast doubt on the assignments. (**F**) An equivalent position in the 1.55 Å resolution **3zp8** structure is assigned as an Na^+^ ion occupying a O_ph_.5O_w_ site. (**G**) A partial view of the *MG_diagnosis* “tab1” output, showing the structural environment of the 17 Mg^2+^ ions assigned in **2quw**, with blue arrows indicating the ions highlighted in panels (**B**) and (**C**).

The hammerhead dimer structures **2quw** and **2qus**, solved at resolutions between 2.2 and 2.4 Å, contain 17 and 16 assigned Mg^2+^ ions, respectively. In **2quw** *Cat Wiz* identified only a single Mg^2+^ with a 2.18 Å distance and sufficient space to accommodate a hexahydrated shell (**Figure 11B**). However, unlike the 2.0 Å resolution Mn^2+^-containing structure **2oeu** (**Table S1**), no clear octahedral water densities were observed. Five of the remaining metal densities in **2quw** could be assigned to hydrated Mg^2+^ ions or to anions. On the other hand, the remaining ten assigned Mg^2+^ ions are poorly positioned and sterically incompatible with Mg^2+^ binding according to *Cat Wiz.* Placement of a hydrated Mg^2+^ ion at these sites would induce severe clashes with neighboring residues (**Figure 11C**).

The absence of well-defined Mg^2+^ binding sites, particularly when contrasted with structures containing transition metals or Na^+^, complicates mechanistic interpretations on currently assigned ion positions. Future investigations of these hammerhead ribozymes may therefore be required to clarify the catalytic mechanisms of hammerhead ribozymes(74,87,88). At present, available hammerhead structures underscore the challenges associated with interpreting solvation shells in low-resolution systems that contain stereochemically unsupported Mg^2+^ assignments which at best can only support speculative interpretations.

Another set of hammerhead structures containing questionable Mg^2+^ assignments include **5dh6**, **5dqk**, and **5eao**. In **5dh6,** the proposed OP2:103 Mg^2+^ binding provides weak support for specific Mg^2+^ coordination as *d*(Mg^2+^…O) ≈2.48 Å exceeds typical Mg^2+^…O distance (**Figure 11D**). Similarly, **5dqk** and **5eao** show densities close to 2.45 Å with no octahedral coordination geometry (**Figure 11E**), suggesting that these features are more consistent with other solvent particles than Mg^2+^ ions. The large number of poorly assigned Mg^2+^ ions in these structures significantly further diminishes confidence in these interpretations. By contrast, the 1.55 Å resolution structure **3zp8**, crystallized in a 1.7 M sodium malonate buffer, displays several clearly defined octahedrally coordinated Na^+^ ions (**Figure 11F**).

In retrospect, early stereochemical validation of these metal-binding sites could have substantially clarified mechanistic interpretations of hammerhead ribozyme. These observations highlight the importance of comprehensive analysis across all available structures of a given system, as well as the need to verify that metal ion assignments conform to fundamental stereochemical principles. A detailed understanding of crystallization buffer composition is equally critical to avoiding misassignments. Overall, *Cat_Wiz* proved highly effective for reevaluating these hammerhead systems.

### Case III: group I *Tetrahymena* ribozyme structures

*Cat_Wiz* was also used to reinspect the solvation shell of different *Tetrahymena* group I ribozyme structures. The first example is an X-ray structure of the 160-nucleotide P4-P6 ribozyme core (PDBid: **1hr2**) solved at 2.25 Å resolution. This fragment is part of the full-length 387-nucleotide ribozyme (89). Its analysis previously led to the description of a well-defined Mg^2+^ core with 42 assigned divalent ions (90). Further, a second set of cryo-EM structures of the full-length *Tetrahymena* ribozyme deposited under PDBids **9cbx** and **9cby** at 2.20 Å and 2.30 Å resolutions, respectively, was analyzed. In these structures, Mg^2+^ ions and water molecules were automatically assigned using a segmentation-guided water and ion modelling protocol called SWIM (22,59,91), and they display slightly different P9.2 stem configurations. Companion structures (**9cbu** and **9cbw**) which contain only the consensus solvent molecules common to **9cbx** and **9cby** were also deposited to the PDB.

*• The P4-P6 mutant solvation shell:* The **1hr2** X-ray structure of this 160-nucleotide group I intron fragment comprises two biological assemblies (89). Chain “A” contains 12 hexacoordinated Mg^2+^ ions that meet our stereochemical criteria and 12 that do not (**Table 3**). Of the validated ions, only four possess a complete hydration shell and establish direct contacts to the RNA. One of these binding sites is of the *mer-*3O_ph_.3O_w_ type (**Figure 12A**). The coordination spheres of three other ions could be adjusted to satisfy the *d*(Mg^2+^…O/N) < 2.3 Å criterion. In addition, eight other hexahydrated Mg^2+^ ions were modelled to form hydrogen bonds with guanine O6/N7 atoms. The hydration shells of the remaining nine ions were not modelled, making it difficult to determine whether they are Mg[H_2_O]_6_^2+^ ions, cacodylate anions, or HEPES fragments based solely on the available density maps. The *Cat_Wiz* analysis confirmed the Mg^2+^ placements for the hydrated ions, while the *MG_detect* tool did not identify any additional inner-sphere Mg^2+^ binding sites, suggesting that all directly bound Mg^2+^ ions in the P4-P6 structure have been accounted for.
*• The full-length **9cbu** and **9cbx** ribozyme solvation shells:* The **9cbu** and **9cbx** structures contain 22 (consensus) and 54 Mg^2+^ ions, respectively. Surprisingly, hydration shells were not modelled for these ions, even though coordinated water molecule densities are apparent for six of these ions. Thus, the positions of these ions could not be validated by *MG_diagnosis*. To complete the coordination analysis of **9cbu**, we examined each of the 22 assigned Mg^2+^ ions individually. Densities for the three water molecules of the *fac/mer-*3O_ph_.3O_w_ metals and the four water molecules of the *cis-*2O_ph_.4O_w_ metals were visible, supporting these Mg^2+^ assignments. Yet at least four Mg^2+^ ions identified in the **1hr2** structure were absent in **9cbu**, despite clear densities supporting their presence (**Figures 12C/D/E** and **Table 3**). More concerningly, 16 Mg^2+^ ions were placed at locations lacking visible octahedral density. Modeling a full hydration shell at these locations result in steric clashes with neighboring residues (**Figure 12F**) suggesting that these sites are unlikely to represent *bona fide* Mg^2+^ binding sites.

**Figure 12.**
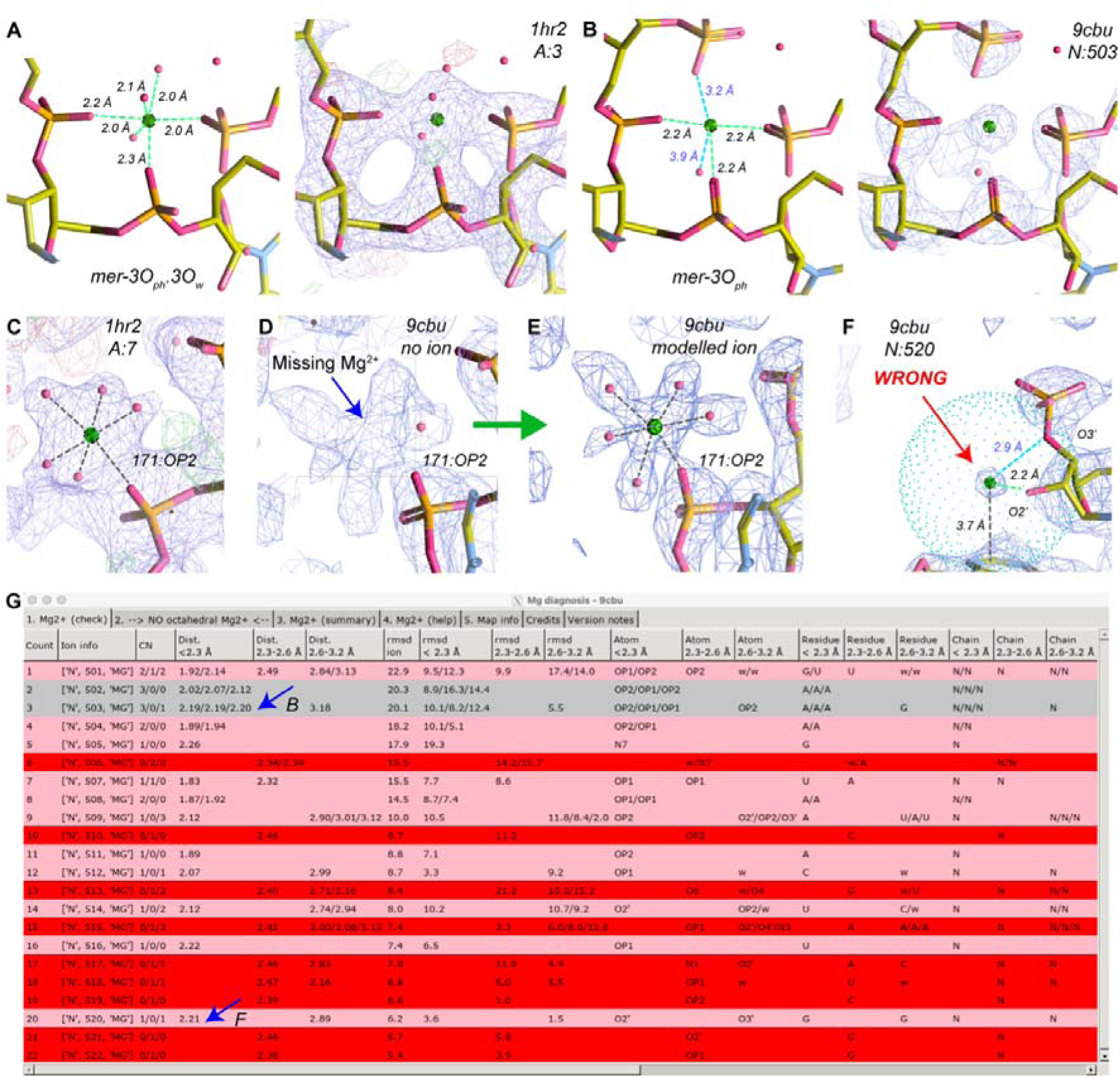
Ion binding assignments in two *Tetrahymena* group I intron structures: (**A**) A *mer-*3O_ph_.3O_w_ binding site in the 2.25 Å resolution **1hr2** X-ray structure (89), and (**B**) the equivalent site modelled in the 2.20 Å resolution **9cbu** cryo-EM structure (22). (**C**) An O_ph_.5O_w_ binding site within an octahedral density pattern in **1hr2**, and (**D-E**) the corresponding site in **9cbu**, which remains unoccupied despite clear octahedral density (**D**) but can be modelled as O_ph_.5O_w_ using *MG_diagnosis* (**E**). (**F**) A 3.6 Å radius sphere, generated by *MG_diagnosis*, emphasizes contacts established by an incorrectly modelled Mg^2+^ ion in **9cbu** positioned too close to an O3’ atom and unrealistically stacking over an adenine nucleobase. (**G**) A partial view of the *MG_diagnosis* “tab1” summarizing the structural environment of the 22 Mg^2+^ ions assigned in **9cbu**, with blue arrows marking the ions shown in panels (**B)** and **(F**).

**Table 3.**
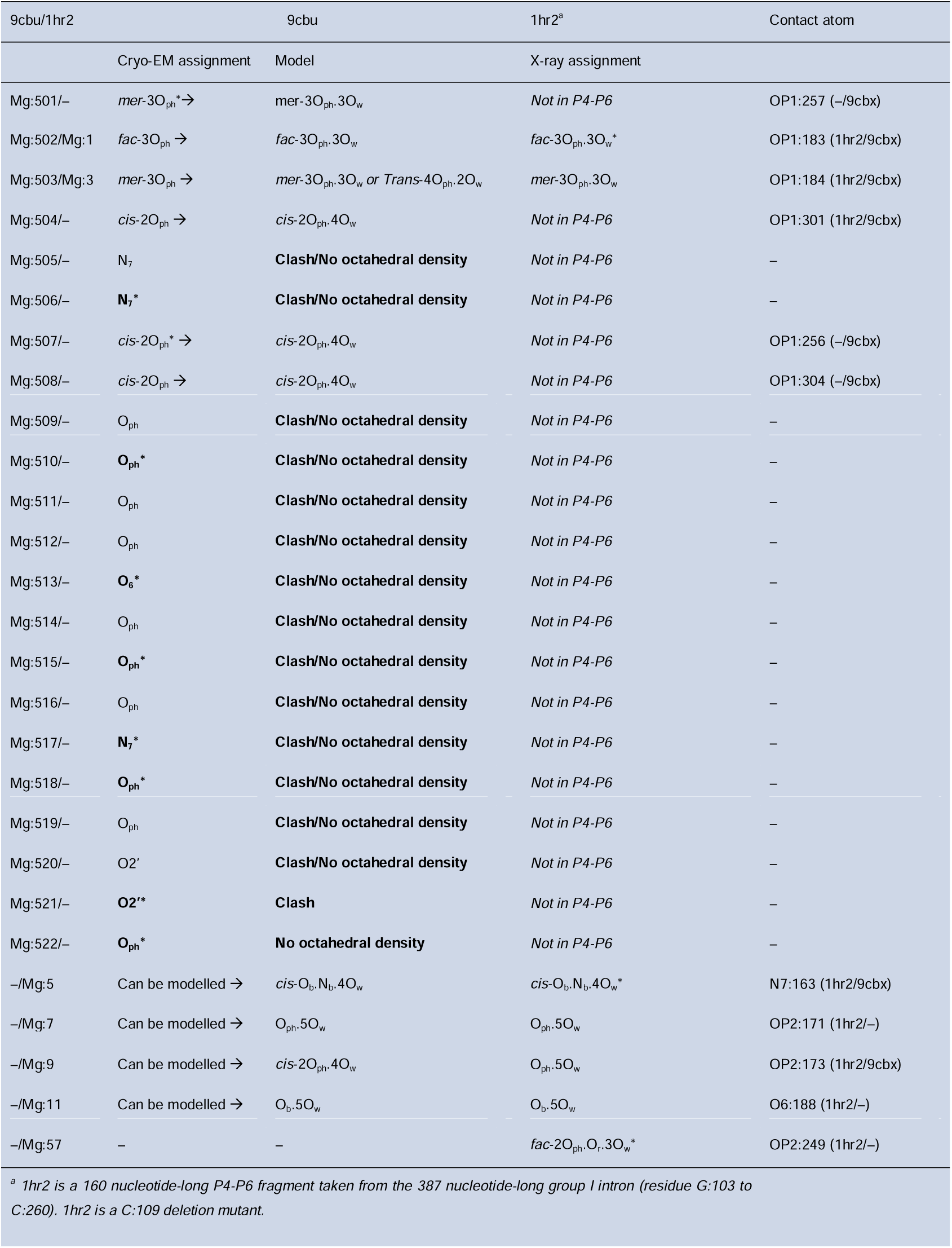
Comparison of Mg^2+^ binding sites in the 9cbu cryo-EM and 1hr2 X-ray *Tetrahymena* group I intron structures. An asterisk indicates the presence of *d*(Mg…O/N) distances in the 2.3–2.6 Å range. In the **1hr2** structure (89), the 10 chain “A” ions (Mg:51 to Mg:64) with no octahedral densities are distant from RNA and at best speculative. These densities should more likely be attributed to other buffer components. The eight assigned Mg[H_2_O]_6_^2+^ chain “A” (Mg:14 to Mg:27) ions are valid, as some of their water molecules form hydrogen bonds with guanine O6/N7 atoms or OP atoms. In **9cbu** (22), 16 out of 22 Mg^2+^ ions display incorrect stereochemistry and are associated with clashes when hydrated. The ions in the consensus **9cbx** structure are shown in the last column.

Regarding non-consensus ions, we found that the ion modelled in **Figures 12C/D/E** is absent in all four parent structures. Furthermore, none of the 32 additionally identified Mg^2+^ ions fit our stereochemical criteria (14). *MG_diagnosis* helped visualize clashes through a sphere of dots, as shown in **Figure 12F**. Our analysis shows that these ions do not display identifiable octahedral densities and are primarily located at sites where a complete hydration sphere induces clashes. The *MG_detect* tool failed to identify any other credible positions for Mg^2+^ ions, despite the presence of numerous unassigned densities.

Taken together, our findings suggest that the *Tetrahymena* group I ribozyme requires only a few precisely positioned Mg^2+^ ions to adopt its active conformation. This analysis demonstrates how *Cat_Wiz* a critical inspection of solvation shells in moderate resolution structures and effectively discriminates between improbable and valid ion assignments. A more comprehensive view of the solvation shell of these group I introns will require higher-resolution structures, preferably obtained using K^+^ rather than Na^+^ containing buffers, as used in most deposited structures.

### Case IV: RNA quaternary structures

- *Exploring metal binding sites in RNA scaffold structures:* Recent advances in cryo-EM have opened new avenues for investigating RNA quaternary assemblies (34). Metal ions were assigned in two such structures: a *Lactobacillus salivarius* ROOL hexamer (PDBid: **9j6y**) and a *Clostridium botulinum* OLE dimer (PDBid: **9lcr**), resolved at 3.25 and 2.6 Å, respectively. At these resolutions, metal hydration shells were not modelled. Noteworthy, in the publication describing these quaternary structures (34), the assigned ions were labelled as “Metals” or (M), except in the supplementary material. In the corresponding PDB file, some of these ions are incorrectly annotated as “Mg” highlighting how automatic database surveys can be compromised by annotations that do not adhere to basic stereochemical criteria. We therefore used these structures to evaluate *Cat_Wiz* performance on low-resolution models and to assess its ability to identify plausible Mg^2+^ binding sites and clamps. In addition, we analyzed a higher-resolution structure of a *Ligi lactobacillus salivarius* ROOL monomer (PDBid: **9m79**), solved at 1.96 Å, for which no ions or water molecules were assigned despite the improved resolution (92).
- *The **9j6y** Lactobacillus salivarius ROOL hexamer (3.25 Å):* In this structure, 56 non-hydrated Mg^2+^ ions were assigned. Of those, 18 form contacts with the RNA in the 2.6–3.2 Å range with coordination distances specific to K^+^ ions. The authors suggested in the supplementary material that a few of the assigned metals form a *“magnesium-clamp”* involving the P13 and P15 secondary structure elements (M6 site). However, *Cat_Wiz* identified *d*(O_ph_…O_ph_) distances of approximately 3.8 Å at this site, which are more consistent with K^+^ binding rather than with Mg^2+^ (**Figure 13A**). The remaining 38 ions, located more than 3.2 Å from RNA atoms, are difficult to assign without additional experimental evidence. While they could relate to hexahydrated Mg^2+^ ions, they may also represent other buffer components. For example, ions placed on the three-fold symmetry axis (M10 site) might correspond to Mg[H_2_O]_6_^2+^ ions given the cavity size. However, the absence of clear octahedral density raises doubts about this assignment (**Figure 13B**).
- *The **9lcr** Clostridium botulinum OLE dimer (2.6 Å):* In this dimer, hydration shells were not modelled for the 28 assigned metals (14 per monomer). Among the 14 ions in monomer “A”, one valid *mer-*3O_ph_.3O_w_ coordination site (M2) was identified. Here, metal coordination distances are below 2.3 Å and the *cis-d*(O_ph_…O_ph_) distances are 2.7/3.1 Å. Four additional unclear octahedral metal sites could correspond to hexahydrated Mg^2+^ ions, while the remaining nine occupy K^+^ sites. These sites, especially M6, M7, M9, M11, M12 and M14, are located near the major groove of G•U pairs which are known to be preferential K^+^ binding sites (3,11,93).

Given the limited resolution of this structure, only remodel a subset of the assigned metals could be remodelled. At the *mer-*3O_ph_.3O_w_ binding site (M2), densities corresponding to three missing water molecules were visible (**Figure 13C**). Using *Coot*, we placed water molecules at their presumed location and reran *MG_diagnosis* followed by positional optimization using the *“Sphere refine with restraints”* option. In a similar manner, we added water to a presumed hexahydrated Mg^2+^ ion at the M1 site and to an M5 ion at a *trans*-2O_ph_.4O_w_ site. This procedure yielded satisfactory hydration sphere model, validating the presence of octahedral coordinated Mg^2+^ ions. A more in-depth remodeling of these structures, including the evaluation of additional orphan densities in the experimental maps is beyond the scope of this paper.

- ***9m79*** *Ligi lactobacillus salivarius ROOL monomer (1.96 Å):* Despite its higher resolution, no ions or water molecules were modelled in this structure (92). Yet *MG_clamp* identified at least three *cis-*2O_ph_.4O_w_ clamps, one *cis-*O_ph_.N_b_.4O_w_ and one relatively well-resolved *fac*-3O_ph_.3O_w_ binding site, demonstrating the usefulness of these ion detection techniques.

**Figure 13.**
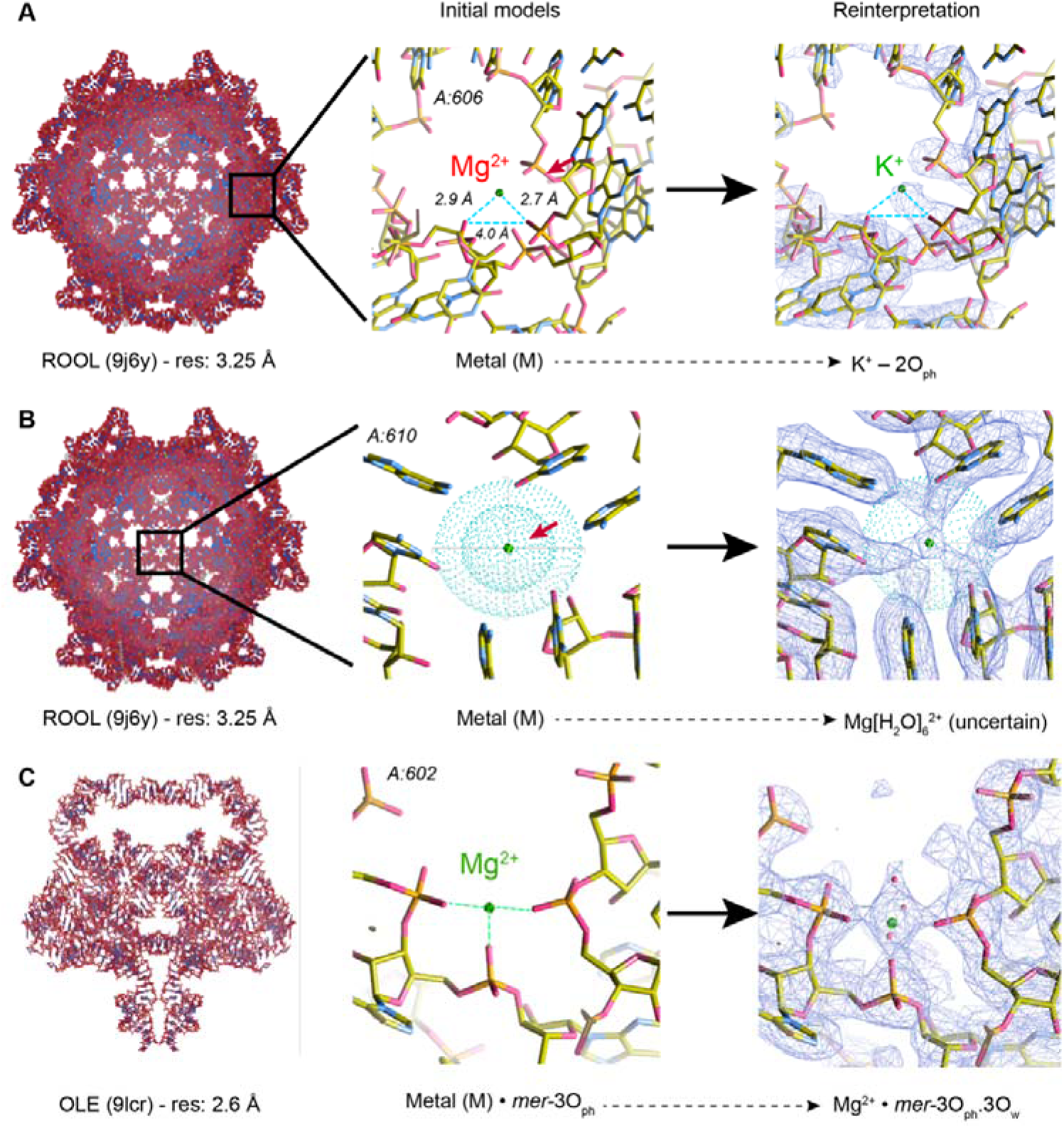
Ion placements in two low and one high-resolution quaternary structures. **(A)** In the **9j6y** structure (34), Mg^2+^ ions with coordination distances in the 2.6–2.9 Å range should be assigned as K^+^ ions. **(B)** This ion is located on a three-fold symmetry axis. The associated non-octahedral densities are difficult to interpret, As visualized by spheres of dots with 2.3 and 3.6 Å radii generated by *MG_diagnosis*, the distances to the neighboring residues allow for the placement of a Mg[H_2_O]_6_^2+^ ion. However, alternative assignments should be considered. **(C)** In this 2.6 Å resolution OLE structure a density pattern near three O_ph_ atoms results in a valid *mer-*3O_ph_.3O_w_ assignment. The three missing water molecules can be modelled without introducing clashes by using *Coot* tools and *Cat_Wiz “Sphere refine with restraints”* options.

Overall, these findings on quaternary assemblies indicate that Mg^2+^ and K^+^ ions in low-resolution structures can in some cases, be differentiated or invalidated based on density patterns and stereochemical considerations. Such prior knowledge-based approaches are essential to limit the propagation of misassignments in structural databases. In these examples, *MG_detect* and *MG_clamp* did not identify additional Mg^2+^ binding sites beyond those discussed. Besides providing a clear survey of the assigned ions, *MG_diagnosis* proved particularly useful for modeling Mg^2+^ hydration shells in these challenging low-resolution structures.

## DISCUSSION

Accurate assignment of Mg^2+^ binding sites remains one of the most significant challenges in RNA structural biology. Misplaced ions are common in the PDB even in high-resolution models where stereochemical constraints are violated or hydration shells are incomplete (11,12,14,94). Such errors and incompleteness lead to inaccurate mechanistic interpretations can mislead functional hypotheses, and introduce noise into databases used for comparative analyses or machine-learning workflows (10,20,21,41–43,95).

*Cat_Wiz* addresses these challenges by integrating stereochemical binding principles into a unified analysis pipeline. By enforcing established coordination distances and geometries derived from the CSD and from a 1.55 Å resolution structure of an *E. coli* ribosome (14), *Cat_Wiz* identifies inconsistent assignments, suggests alternative interpretations and highlights regions where experimental density or backbone modeling may be inadequate. Herein, we demonstrated applications to ribosomes, hammerhead ribozymes, group I introns, and RNA quaternary assemblies, illustrating how *Cat_Wiz* can recover incomplete ionic hydration shells, correct misassigned cations, and annotate ion motifs such as Mg^2+^ clamps. Importantly, these corrections can be achieved within hours rather than days of manual examination. *Cat_Wiz* is applicable to both X-ray and cryo-EM structures across a broad resolution range and can be used for nucleic acids as well as proteins.

While *Cat_Wiz* provides valuable insights into RNA structural biology it shares limitations common to all structure-based approaches. At resolutions poorer than ≈2.5 Å, ion coordination becomes increasingly difficult to distinguish and errors in RNA backbone modeling may propagate into ion placement. To mitigate these issues, *Cat_Wiz* provides restrained refinement options that enforce realistic Mg^2+^ coordination distances while preserving structural flexibility. These restraints are conceptually analogous to those routinely applied to nucleotides and amino acids and aim to generate chemically acceptable structural models (96–98).

Buffer composition presents a second challenge. Non-physiological intracellular cations (99–101) such as NH_4_^+^, Na^+^, and anions including cacodylate and sulfate (82,83) are frequently present in experimental buffers. Their coordination distances can overlap with those of Mg^2+^, complicating density interpretation and increasing the likelihood of misassignment. *Cat_Wiz* explicitly highlights cases where such ions have been misinterpreted as Mg^2+^, as illustrated here for hammerhead ribozymes. These findings strengthen the importance of using simplified and physiologically relevant buffers containing primarily Mg^2+^ and K⁺, to warrant better density map interpretability and ensure physiologically meaningful assignments. Notably, the use of *“simplified”* buffers does not preclude the determination of high-resolution structures and is compatible with *in vitro* autonomous ribosome biogenesis (19,102). In the future, more systematic inclusion of polyamines in experimental buffers may further advance our understanding of biological solvation shells (62,103–106). Although chloride (Cl^−^) is the most abundant intracellular anion (3,82,83), it is often absent from modern experimental buffers and replaced by other anionic components including acetate, malonate, HEPES or TRIS molecules. Such substitutions may perturb RNA solvation shells in ways that remain poorly understood.

A third concern relates to the deposition and annotation of structural data (15,18,107,108). Misassigned ions lacking appropriate annotations persist in the PDB, despite repeated identifications of such errors (15,32). Greater and more consistent user of the CAVEAT remark would help flag known issues in deposited PDB structures (18). Such inaccuracies not only mislead newcomers to the field but also comprise automated surveys and machine-learning training datasets. Herein, we documented how such issues affect hammerhead and recent group I intron ribozyme structures potentially jeopardizing collective efforts to understand and model their ionic structure and catalytic properties (22). Broader adaption of annotation tools such as *Cat_Wiz*, together with improved PDB-level flagging, would significantly enhance the overall quality of publicly available structural models. As a side note, the PDB currently lacks specific residue codes for anionic U(-) and G(-) nucleotides recently identified in bacterial ribosomes (e.g., **8b0x**) despite the existence of dedicated codes for protonated A(+) and C(+) residues (“AP7” and “CH”, respectively) (14,65). Addressing this limitation would further improve ion and charge-state annotation.

*Cat_Wiz* also highlights the inherent limits of automation. Alternate nucleotide conformations, ambiguous solvent densities, and unusual motifs such as metal-mediated base pairs or diffuse ion populations, require expert visual inspection prior to publication. *Cat_Wiz* is designed to flag these instances, yet it does not aim at automatically resolving them, as careful human interpretation remains essential for understanding and documenting these features. Even with *Cat_Wiz*-assisted inspection, analyses of poorly resolved regions may still raise more questions than definitive answers.

Finally, while machine-learning ion identification approaches are gaining popularity (20,109,110), current methods struggle to correct stereochemical errors and misassignments already present in existing models. Similarly, MD simulations that lack polarization and charge transfer effects must be interpreted cautiously when modelling Mg^2+^ ions (22,111,112). By contrast, *Cat_Wiz* provides a crucial bridge between experimental data and modelling efforts. By incorporating rigorous stereochemical principles and generating high-quality ion assignments, it supplies curated datasets that can improve the accuracy and robustness of next generation AI tools and molecular dynamics force fields (113–117).

To summarize, *Cat_Wiz* offers a rapid, and stereochemistry-guided framework for diagnosing, annotating and correcting Mg²⁺ assignments in biomolecular structures. By improving the fidelity of solvation-shell models and producing carefully curated datasets, *Cat_Wiz* benefits experimentalists seeking accurate mechanistic interpretations and computational scientists developing predictive models of ion-RNA interactions. Its integration into *Coot* makes it broadly accessible, positioning *Cat_Wiz* as a practical, fast, and transparent tool supplementing both manual refinement and emerging autonomous structural biology approaches.

### Disclaimer

We made every effort to design *Cat_Wiz* rigorously and to assign Mg^2+^ ions in the **8b0x/9q87** structure as accurately as possible. The quality of these assignments necessarily depends on structural resolution, preceding refinement steps, and the state of prior knowledge available at the time of structure deposition, particularly when disentangling complex or unforeseen solvation patterns. This remains an ongoing process. Accordingly, we do not claim: *i)* that all Mg^2+^ ions were identified, *ii)* that particularly complex or poorly resolved solvation patterns did not occasionally lead to ambiguous interpretations, or *iii)* that all reported coordination patterns are free of modeling imperfections. However, our analyses were conducted using the best available structural data and current stereochemical knowledge. We are confident that all major Mg^2+^ binding sites in the **8b0x** ribosomal structure were identified and that the most relevant Mg^2+^ binding types across the RNA systems examined were captured, enabling a near-complete and internally consistent annotation framework.

## AVAILABILITY

The scripts are deposited to Zenodo as password protected files at the following address: https://zenodo.org/records/18144017.

## ACCESSION NUMBERS

Atomic coordinates and structure factors for the reported crystal structures have been deposited to the PDB under accession number **9q87** (version 3).

## SUPPLEMENTARY DATA

Supplementary Data are available at NAR online.

## ACKNOWLEDGEMENT

The authors thank Prof. Eric Westhof and Dr. Eric Ennifar for helpful discussions, Dr. Ahmad Jomaa and Dr. Emir Maldosevic for providing info on cryo-EM structures, Dr. Paul Emsley and Dr. Lucrezia Catapano for help and advice on *Coot* functionalities, as well as Dr Aliza Fedorenko, Dr Anat Bashan and Dr Jiro Kondo for discussions on the possible presence of copper as contaminant in the experimental process. We also thank Prof. Giovanni Bussi that organized a CECAM meeting in Paris where P.A and S.K. first met and started this collaborative research project.

## Author contributions

Nawavi Naleem (Conceptualization [equal], Methodology [equal], Software [lead], Validation [equal], Visualization [equal], [Writing - review & editing [supporting]), Anja Henning-Knechtel (Conceptualization [supporting], Methodology [supporting], Writing - review & editing [supporting]), Serdal Kirmizialtin (Conceptualization [supporting], Funding acquisition [lead], Methodology [supporting], Project administration [lead], Resources [lead], Software [supporting], Supervision [supporting], Writing - review & editing [equal]), Pascal Auffinger (Conceptualization [lead], Methodology [equal], Project administration [supporting], Software [supporting], Supervision [lead], Validation [equal], Visualization [equal], Writing - original draft [lead], Writing - review & editing [equal]). d

