## Supplemental 1 for "Cat_Wiz: A stereochemistry-guided toolkit for locating, diagnosing and annotating Mg^2+^ ions in RNA structures"

**Table of content:**

1) Extra distance restraints definition.
2) Nomenclature issues regarding the presence of O1P/O2P in some modified residues.
3) Command to replace the “_atom_site.label_seq_id” by the “_atom_site.auth_seq_id” column in a cif file.
4) Avoid “TER” lines in PDB and cif files.
5) **Table S1:** List of the 25 *Hammerhead* ribozyme X-ray structures deposited to the PDB.
6) **Figure S1:** Modelisation of a copper cation in a density bridging two residues of r-protein L27.
7) **References:**

**I. Extra distance restraints definition=**

All the *Coot* commands can be found at this and related web sites.

For instance, see: <https://www2.mrc-lmb.cam.ac.uk/personal/pemsley/coot/docs/html/files.html>

**add_extra_bond_restraint** (imol, chain_id_1, res_no_1, ins_code_1, atom_name_1, alt_conf_1, chain_id_2, res_no_2, ins_code_2, atom_name_2, alt_conf_2, bond_dist, esd)

As an example, the B:205:OD1 atom is at 3.19 Å from 9:3969:MG. This short distance prevents validation of this relatively poorly resolved site. An appropriate restraint line would be:

**add_extra_bond_restraint** (0, ‘B’, 205, ‘’, ‘OD1’, ‘’, ‘9’, 3969, ‘’, ‘MG’, ‘’, 3.4, 0.001) and then use “*sphere refine with restraint*” option. Afterwards, eventually use *“delete_all_extra_restraints*”.

**II. Nomenclature issues regarding the presence of O1P/O2P in some modified residues.**

Some residues like 6MZ still are defined with O1P/O2P instead of OP1/OP2. It is important to pay attention to such details.

**III. Command to replace the “_atom_site.label_seq_id” by the “_atom_site.auth_seq_id” column in a cif file.**

It is sometimes necessary to swap these columns in the *Coot* cif files. The command is:

awk ' {t = $23; $9 = t; print; } ' input.cif > output.cif

**IV. Avoid “TER” lines in PDB and cif files.**

The presence of lines starting with “TER” in PDB and cif files can cause misfunctions of the *Cat_Wiz* scripts.

**Table S1.** **List of the 25 *Hammerhead* ribozyme X-ray structures deposited to the PDB.** The structures with assigned Mg^2+^ ions are in bold. CAC is the PDB code for cacodylate anions (AsO_2_(CH_3_)_2_^-^). The crystallization buffer compositions were derived from the original publications and are only indicative.

|  | PDB | Date | Res.(Å) | Ions | Nucl. | Buffer components | Ref |
| --- | --- | --- | --- | --- | --- | --- | --- |
| 01. | 1hmh ^a^ | 1995 | 2.6 | – | 34+13 | 2.4M Li_2_SO_4_; 10mM MgCl_2_; CAC; |  |
| 02. | 1mme | 1996 | 3.1 | – | 25+16 | 100mM NH_4_OAc; 10mM Mg(OAc)_2_; CAC; |  |
| 03. | 299d | 1997 | 3.0 | – | 25+16 | 1.8M Li_2_SO_4_; 10mM MgSO_4_; CAC; |  |
| 04. | 300d | 1997 | 3.0 | 6 Mn^2+^ | 25+16 | ^b^ |  |
| **05.** | **301d** | **1997** | **3.0** | **5 Mg^2+^** | **25+16** | 1.8M Li_2_SO_4_; 10mM MgSO_4_; CAC; |  |
| 06. | 359d | 1997 | 2.9 | 5 Tb^3+^ | 25+16 | ^b^ |  |
| 07. | 379d ^a^ | 1998 | 3.1 | 8 Co^2+^ | 25+16 | ^b^ |  |
| 08. | 488d ^a^ | 2000 | 3.1 | 8 Cd^2+^ | 20+16+25+5 | ^b^ |  |
| 09. | 1nyi ^a^ | 2004 | 2.85 | 4 Co^2+^ | 24+16 | ^b^ |  |
| 10. | 1q29 ^a^ | 2004 | 3.0 | 5 Co^2+^ | 24+17 | ^b^ |  |
| 11. | 2oeu | 2007 | 2.0 | 5 Mn^2+^ | 43+20 | ^b^ |  |
| **12.** | **2quw** | **2008** | **2.2** | **17 Mg^2+^** | **57+12** | 28mM MgCl_2_, CAC; see |  |
| **13.** | **2qus** | **2008** | **2.4** | **16 Mg^2+^** | **69** | 28mM MgCl_2_, CAC; see |  |
| 14. | 3zd4 | 2012 | 2.2 | – | 43+20 | 1.7M Na-malonate; 10mM MgCl_2_; |  |
| 15. | 3zd5 | 2012 | 2.2 | – | 43+20 | 0.5M (NH_4_)_2_SO_4_; 1mM MgCl_2_; |  |
| 16. | 3zp8 | 2013 | 1.55 | 16 Na^+^ | 43+20 | 1.7M Na-malonate; 10mM MgCl_2_; |  |
| **17.** | **5dh6** | **2015** | **2.78** | **5 Mg^2+^** | **48+20** | 0.9 M KCl; 10mM MgCl_2_; CAC; |  |
| 18. | 5dh7 | 2015 | 3.06 | 11 Mn^2+^ | 48+20 | ^a,b^ |  |
| 19. | 5dh8 | 2015 | 3.30 | 13 Zn^2+^ | 48+20 | ^a,b^ |  |
| 20. | 5di2 | 2015 | 2.99 | 7 Mn^2+^ | 48+20 | ^a,b^ |  |
| 21. | 5di4 | 2015 | 2.95 | 11 Mn^2+^ | 48+20 | ^a,b^ |  |
| **22.** | **5dqk** | **2016** | **2.71** | **1 Mg^2+^ + 4 K^+^** | **48+20** | 50mM Mg(OAc)_2_; CAC; |  |
| **23.** | **5eao** | **2016** | **2.99** | **8 Mg^2+^** | **48+20** | 0.4M KCl; 50mM Mg(OAc)_2_; CAC; |  |
| 24. | 5eaq | 2016 | 3.2 | 6 Mn^2+^ | 48+20 | ^a,b^ |  |
| 25. | 8ydc | 2024 | 2.89 | – | 58+14 | 2.5M (NH_4_)_2_SO_4_; 10mM Mg(OAc)_2_; |  |
| ^a^ Density maps were not deposited to the PDB.  ^b^ For the buffer composition of the structures crystallized in the absence of Mg^2+^ ions,  please refer to the original publications. | | | | | | | |

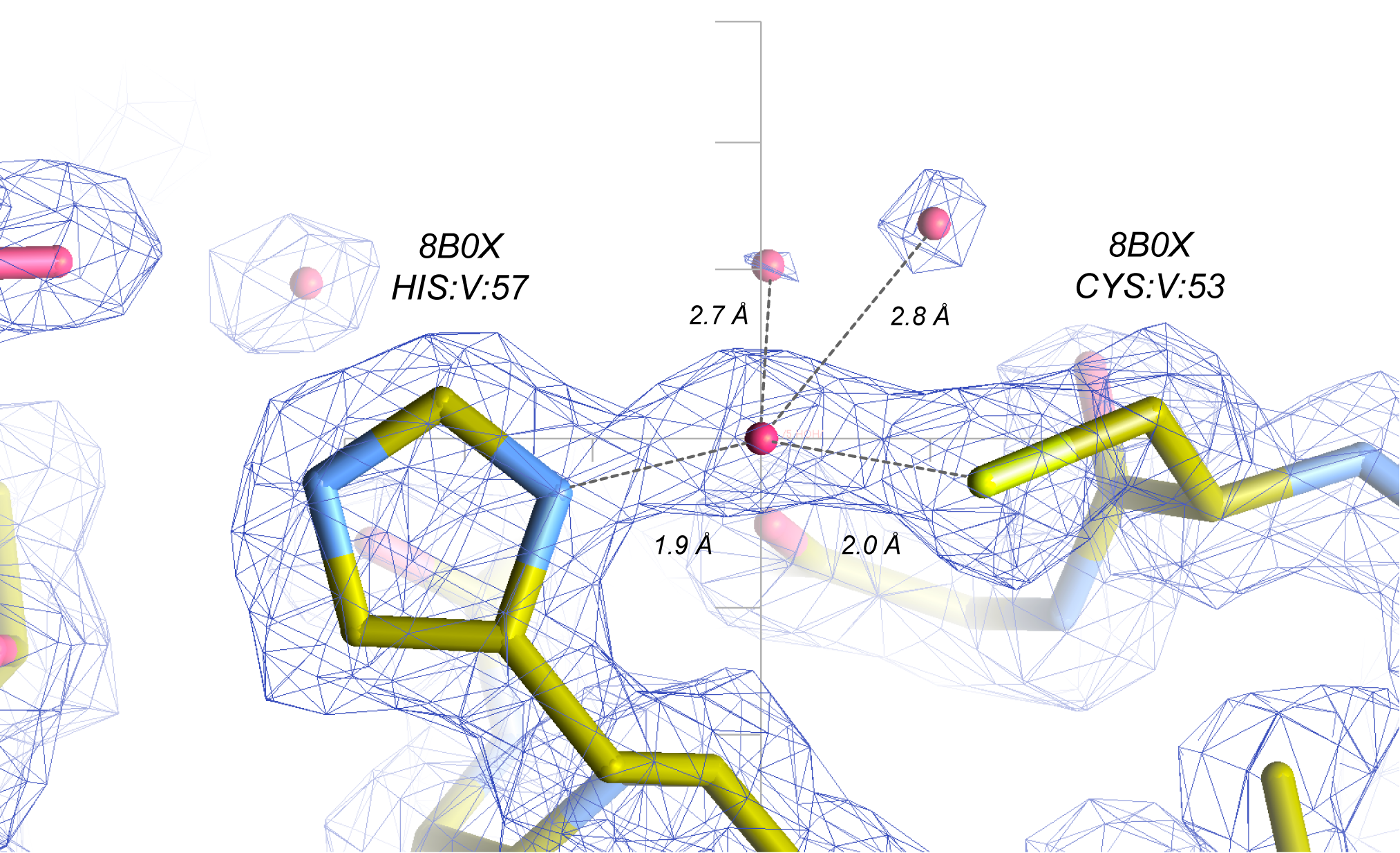

**Figure S1. Modelisation of a copper cation in a density bridging two residues of r-protein L27.** This ion is modelled in **8b0x**.
